# Systems-level transcriptional regulation of *Caenorhabditis elegans* metabolism

**DOI:** 10.1101/2022.11.08.515576

**Authors:** Shivani Nanda, Marc-Antoine Jacques, Wen Wang, Chad L Myers, L. Safak Yilmaz, Albertha JM Walhout

## Abstract

Metabolism is precisely controlled to ensure organismal development and homeostasis. Several mechanisms regulate metabolism, including allosteric control and transcriptional regulation of metabolic enzymes and transporters. So far, metabolism regulation has mostly been described for individual genes and pathways, and the extent of transcriptional regulation of the entire metabolic network remains largely unknown. Here, we find that more than three-quarters of all metabolic genes are transcriptionally regulated in the nematode *Caenorhabditis elegans*. We find that many annotated metabolic pathways are coexpressed, and we use gene expression data and the iCEL1314 metabolic network model to define coregulated sub-pathways in an unbiased manner. Using a large gene expression compendium, we determine the conditions where sub-pathways exhibit strong coexpression. Finally, we develop ‘WormClust’, a web application that enables a gene-by-gene query of genes to view their association with metabolic (sub)-pathways. Overall, this study sheds light on the ubiquity of transcriptional regulation of metabolism and provides a blueprint for similar studies in other organisms, including humans.

## Introduction

All organisms regulate their metabolism during development and to maintain homeostasis under fluctuating dietary and environmental conditions. In humans, failure to maintain homeostasis can lead to a variety of metabolic disorders such as inborn errors in human metabolism, obesity, hypertension, and diabetes (DeBerardinis and Thompson, 2012; Sharma et al., 2008). Metabolism can be regulated through different mechanisms. One well-known mechanism is allostery, a fast-acting mechanism where metabolites directly modulate enzyme activity. For instance, the enzyme phosphofructokinase, which regulates the conversion of fructose 6-phosphate to fructose 1,6-biphosphate, is allosterically regulated during glycolysis. This reaction is coupled to ATP hydrolysis where ATP binding to phosphofructokinase inhibits enzyme activity by decreasing its affinity for fructose 6-phosphate, while conversion to AMP reverses the inhibitory effect and increases the activity of the enzyme (Blangy et al., 1968; Schirmer and Evans, 1990). Metabolism can also be regulated transcriptionally by activating or repressing the expression of genes encoding metabolic enzymes or transporters. This mechanism is relatively slow and allows the organism to adapt to changing cellular or environmental conditions. Well-known examples of the transcriptional regulation of metabolism include induction of the lac operon in *E. coli* in response to a switch from glucose to lactose as a carbon source (Gilbert and Muller-Hill, 1966; Jacob and Monod, 1961); the Leloir pathway in yeast which is transcriptionally activated by galactose (Caputto et al., 1949; Hopper et al., 1978); and mammalian cholesterol biosynthesis genes, which are activated by the transcription factor (TF) SREBP when cholesterol levels are low (Brown and Goldstein, 1997; DeBose-Boyd and Ye, 2018). Another example of transcriptional rewiring of metabolism involves propionate degradation in the nematode *Caenorhabditis elegans*. Like humans, the nematode *Caenorhabditis elegans* utilizes a vitamin B12-dependent pathway to breakdown this short-chain fatty acid. When dietary vitamin B12 is low, propionate metabolism is transcriptionally rewired to an alternative degradation pathway referred to as the propionate shunt, thereby preventing toxic propionate accumulation (Bulcha et al., 2019; Watson et al., 2014; Watson et al., 2016). While transcriptional regulation has been studied for individual genes and enzymes, the extent to which overall metabolic activity is under transcriptional control remains unclear due to lack of systems-level studies.

*C. elegans* is an excellent multicellular model to study the transcriptional regulation of metabolism at a systems level: its fixed lineage of 959 somatic cells was fully described (Sulston and Horvitz, 1977), its metabolism shows extensive conservation with human metabolism (Lai et al., 2000; Shaye and Greenwald, 2011), many gene expression datasets are available, and a genome-scale metabolic network model (MNM) has been reconstructed (Yilmaz and Walhout, 2016). The most up-to-date MNM, iCEL1314, contains 907 metabolites, 2,230 reactions and 1,314 genes (Yilmaz et al., 2020). By using flux balance analysis (FBA), iCEL1314 can be used to gain insight into the metabolic state of *C. elegans* during different nutritional conditions or in different tissues. An additional set of metabolic genes has been predicted based on homology with known metabolic enzymes in other organisms or based on the presence of domains found in metabolic enzymes (Bhattacharya et al., 2022; Yilmaz and Walhout, 2016). However, these genes have not been incorporated into the iCEL1314 model because they are not yet annotated to known metabolic reactions or because the reactions they are annotated to cannot yet be connected to the model.

Here, we investigated the extent of transcriptional regulation of *C. elegans* metabolism. We developed a computational pipeline to identify genes of which the corresponding mRNA varies significantly during development, in different tissues, and across a gene expression compendium consisting of different conditions. Using both a supervised and an unsupervised method, we identified coexpressed metabolic pathways and sub-pathways. Overall, we found that more than three-quarters of metabolic genes exhibit variation in expression, which is comparable to the proportion in non-metabolic genes. Further, we found that most annotated metabolic pathways contain genes that are significantly coexpressed. With a custom-made unsupervised method, we identified clusters of genes that define coexpressed sub-pathways or combinations of sub-pathways that likely form functional metabolic units. We extracted conditions where coexpressed clusters of genes are coordinately activated or repressed, revealing how these clusters may contribute to metabolic homeostasis. We developed a web application we named ‘WormClust’ that is available on WormFlux website (Yilmaz and Walhout, 2016). WormClust enables querying of *C. elegans* metabolic genes including transporters, and transcription factors to identify metabolic (sub-)pathways with which these genes are coexpressed. Altogether, our findings show that transcriptional regulation of metabolic genes and pathways is ubiquitous in *C. elegans,* and our analyses and tools provide a platform for similar studies in other organisms, including humans.

## Results

### More than Three-Quarter of Metabolic Genes are Transcriptionally Regulated

mRNA levels are determined by a combination of synthesis and degradation. Here, we used variation in mRNA levels as a first approximation for transcriptional regulation. To this end, we evaluated the expression of metabolic genes during development, in different tissues, and under different conditions to identify metabolic genes that are highly variant and therefore likely transcriptionally regulated. We used all annotated metabolic genes (Yilmaz and Walhout, 2016) and grouped them into four classes based on current annotation (**Table EV1**): Class A, iCEL1314 genes (N=1,308; six genes have been removed because they are annotated as pseudogenes in WormBase WS282, see **Methods**); class B, genes annotated to reactions that cannot yet be connected to the iCEL1314 model (N=192); class C, genes encoding proteins with homology to metabolic enzymes in other organisms (N=860); and class D, genes encoding proteins with a domain found in known metabolic enzymes (N=132). Hereafter, we refer to the 1,308 genes in class A as ‘iCEL1314 genes’ and the remaining 1,184 as ‘other metabolic genes’.

We first identified metabolic genes that vary in expression during development by using a high-quality post-embryonic time-resolved RNA-seq dataset, hereafter referred to as the ‘development dataset’ (**Fig 1A, Fig EV1A, Table EV2**). Briefly, this dataset contains expression profiles of stage-synchronized animals that were collected every two hours after hatching for 48 hours at 20° C. In the original paper, genes were grouped into 12 clusters based on similarity in developmental expression profiles (Kim et al., 2013). One of these clusters contains 5,045 genes, including 995 metabolic genes, with relatively invariant temporal expressions. We will refer to this cluster as the ‘flat cluster’. However, although the expression levels of most of the flat cluster genes are relatively stable during development, we noticed that some did exhibit considerable variation. Additionally, many invariant genes from other clusters were not included in the flat cluster. Therefore, we used an unbiased statistical method to stringently define variation in developmental gene expression. Briefly, we first defined a flat reference by taking the mean +/-standard deviation of normalized expression values of the flat cluster genes over time (**Fig EV1A**). We then developed a statistical measure called variation score (VS) to calculate the deviation of a gene’s expression profile from this flat reference over time (**Fig EV1A**). To define highly variable genes, we empirically established a conservative VS threshold value of 0.169 based on the distribution of VS between flat genes and all other genes, such that 97% of flat genes were not annotated as variant (**Fig EV1B,** see details in **Methods**). After excluding 3,552 lowly expressed genes, including 213 metabolic genes, we found that 754 metabolic genes (31.4%) had a VS greater than this threshold value and were thus annotated as highly variant, whereas 98 metabolic genes (4%) had a VS equal to zero and were annotated as invariant (**Fig 1B**). The remaining 1,332 metabolic genes with VS greater than zero but less than the threshold were annotated as moderately variant (**Fig 1B, Fig EV1C**). About a quarter of iCEL1314 genes (329, or 26%) are highly variant, which is lower than the proportion of other metabolic genes (37%) and non-metabolic genes (41%) (**Fig EV1C-D, Table EV2**). The percentage of highly variant metabolic genes is lower than that of non-metabolic genes across most VS thresholds (**Fig 1C**).

**Figure 1.**
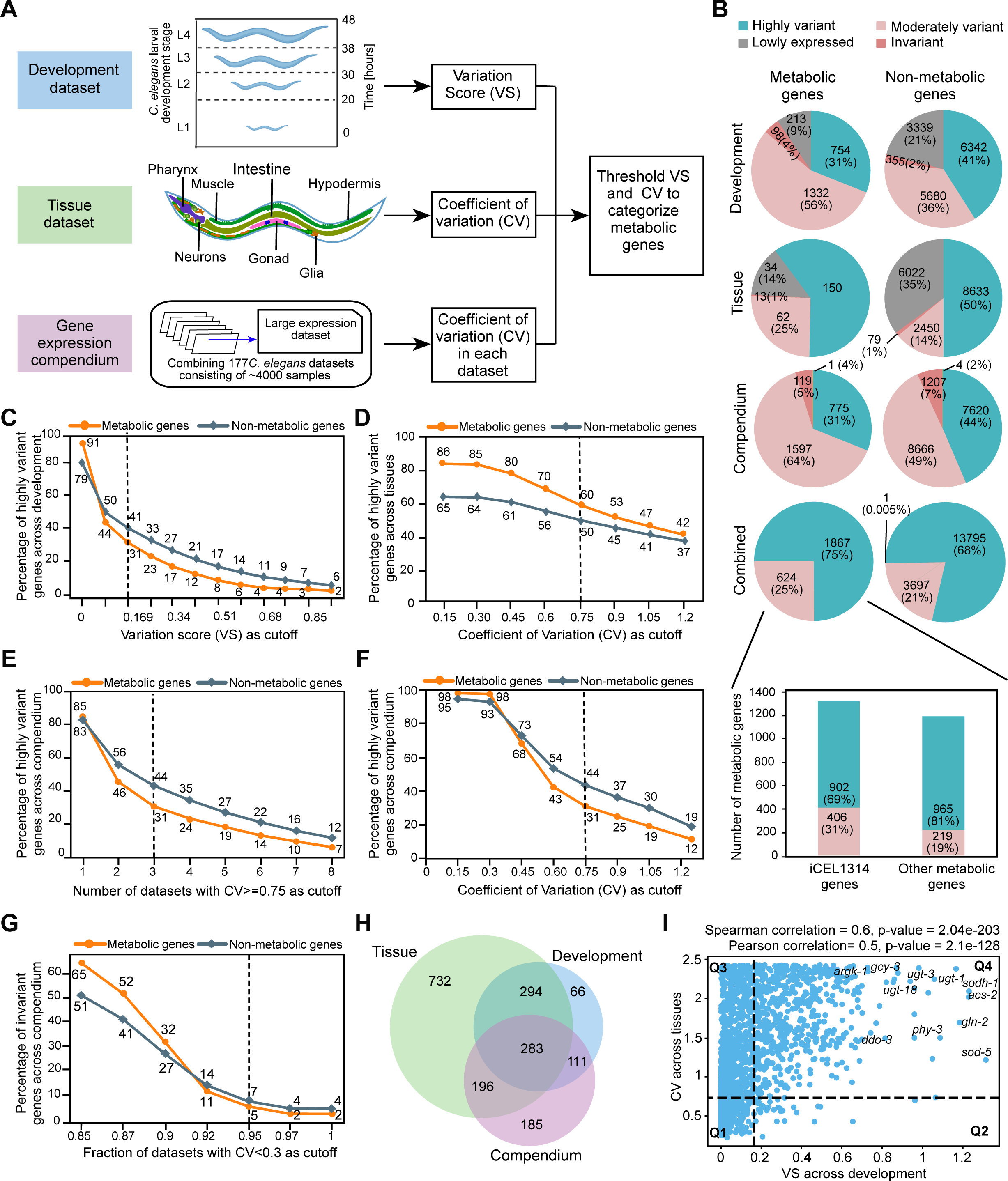
Analysis of Metabolic Gene Expression during Development, in Different Tissues and in a Gene Expression Compendium. (A) Computational pipeline to identify *C. elegans* metabolic genes that change in expression during development, across tissues, and compendium of multiple conditions. Statistically significant differences in gene expression were calculated in the developmental dataset using a variation score (VS), in the tissue dataset using coefficient of variation (CV) and by number of datasets with CV ≥ 0.75 in the compendium (collection of 177 datasets). (B) Pie charts of metabolic and non-metabolic gene expression in the three different datasets: development, tissue, and compendium separately and combined. Bar graph shows metabolic genes in iCEL1314 and other (predicted) metabolic genes. (C) Comparison of percentage of highly variant metabolic versus non-metabolic genes at different VS thresholds. (D) Comparison of percentage of highly variant metabolic versus non-metabolic genes at different CV thresholds. (E) Comparison of percentage of highly variant metabolic versus non-metabolic genes at different cutoffs of number of datasets with high CV (≥ 0.75). (F) Comparison of percentage of invariant metabolic versus non-metabolic genes at different cutoffs of the fraction of datasets with low CV (<0.3). (G) Comparison of percentage of highly variant metabolic versus non-metabolic genes at different CV cutoffs in at least three datasets in the compendium. (H) Venn diagram of highly variant metabolic genes in the different datasets. (I) Scatter plot of VS (development) versus CV (tissue) of metabolic genes. The plot is divided into four quadrants: Q1 with moderate/low VS and moderate/low CV; Q2 with high VS and moderate/low CV; Q3 with moderate/ low VS and high CV; and Q4 with high VS and high CV. The Pearson and Spearman correlation coefficients and the corresponding p-values are indicated.

To identify metabolic genes that exhibit differential expression across tissues, we selected a high-quality single-cell RNA sequencing dataset that measured gene expression during L2 stage of *C. elegans* across seven major tissues: body wall muscle, glia, gonad, hypodermis, intestine, neurons, and pharynx (Cao et al., 2017) (**Fig 1A, Fig EV2A, Table EV3**). This dataset is hereafter referred to as the ‘tissue dataset’. Unlike the development dataset, the tissue dataset does not have a defined cluster of invariant genes. Therefore, we used the statistical measure Coefficient of Variation (CV) to identify variation in gene expression across the seven tissues (**Fig EV2A**). To set a CV threshold for selecting genes that are differentially expressed in distinct tissues, we visually inspected genes with CV values from 0.15 to 1.2 (**Figs EV2A and EV2B**). We previously found that the five genes comprising the propionate shunt are differentially expressed in different tissues (Watson et al., 2016; Yilmaz et al., 2020), and each of these genes had a CV greater than 0.75 (**Fig EV2C**). We, therefore, selected a CV cutoff of 0.75 as a conservative threshold to annotate highly variant genes across tissues. Further, we classified genes with a CV less than 0.75 but greater than or equal to 0.3 as moderately variant, and genes with a CV less than 0.3 as invariant (**Fig EV2D**, see **Figs EV2B and EV2C** for examples). A total of 6,370 genes, including 348 metabolic genes, were not included in this analysis because they are expressed at low levels (Yilmaz et al., 2020)(details in **Methods**). Using these thresholds, we identified ∼60% and ∼25% of metabolic genes as highly and moderately variant, respectively. These include 781 highly variant and 405 moderately variant iCEL1314 genes (**Figs 1B and EV2E**). A very small number of metabolic genes (13, or 1%) were invariant across tissues. These results show that at least twice as many metabolic genes are variant and, therefore, likely transcriptionally regulated in different tissues than during development (**Fig 1B**). In contrast to development, the percentage of metabolic genes that are highly variant across tissues at any CV cutoff is greater than non-metabolic genes **(Fig 1D)**, indicating that metabolic processes exhibit a relatively high level of tissue specificity.

To evaluate metabolic gene expression more broadly, we combined 177 expression profiling datasets into an expression compendium, an earlier version of which we have used to study TF paralogs (Reece-Hoyes et al., 2013) (**Figs 1A and EV3A, Tables EV4 and EV5, see Methods**). To be consistent with the approach used with the tissue dataset, and to be conservative in our assessment, we selected a CV threshold of 0.75 and required that highly variant genes had a CV greater than or equal to this threshold in at least three datasets. Genes that showed CV<0.3 in at least 95% of the datasets were labeled as invariant, and genes that fit into neither category were annotated as moderately variant (**Fig EV3A**). We further assumed that genes that are not present in a dataset are lowly expressed, and we removed these from our analysis. Using these criteria, we found that 775 of the 2,492 metabolic genes (∼31%), including 284 iCEL1314 genes, are highly variant (**Figs 1B and EV3B**). This is significantly lower than the percentage of highly variant non-metabolic genes (44%, **Fig 1B**). This difference holds true for different cutoffs of the number of datasets showing high variation (**Fig 1E**), and across different CV thresholds (**Fig 1F**). However, the percentage of invariant genes is similar between metabolic and non-metabolic genes using different CV cutoffs (**Fig 1G**).

To evaluate the extent of transcriptional regulation in different contexts, we compared highly variant genes in development, tissue, and compendium and found that a total of 1,867 metabolic genes are highly variant in some context, 283 of which are highly variant across all three datasets (**Fig 1H**). Overall, metabolic gene expression showed higher variation across space (tissues) than time (development). To directly compare metabolic gene expression in tissues and development, we plotted VS values of metabolic genes across development versus CV values across tissues and found that these two parameters are moderately correlated (**Fig 1I**). We divided the scatter plot into four quadrants: Q1 with moderate/low developmental variation and moderate/low tissue variation, Q2 with high developmental variation and moderate/low tissue variation, Q3 with moderate/low developmental variation and high tissue variation, and Q4 with high developmental variation and high tissue variation (**Table EV6**).

We performed pathway enrichment analysis (PEA) on metabolic genes for each quadrant using the tool provided on the WormFlux website (Yilmaz and Walhout, 2016). Q1 consists of 595 metabolic genes, including 385 iCEL1314 genes that are enriched in several metabolic pathways, such as the electron transport chain (ETC), aminoacyl-tRNA biosynthesis, the tricarboxylic acid (TCA) cycle, the pentose phosphate pathway and glycolysis/ gluconeogenesis (**Appendi**x **Fig S1**). Q2 consists of only 176 genes, including 68 iCEL1314 genes that are enriched in sulfur, cysteine, and methionine metabolism (**Appendix Fig S1**). The 891 genes in Q3 include 504 iCEL1314 genes that are highly enriched in lipid metabolism. Notably, genes involved in peroxisomal fatty acid (FA) metabolism vary more in expression than mitochondrial FA degradation (**Appendix Fig S1**). Finally, the 577 genes in Q4 include 261 iCEL1314 genes which are enriched in UGT (UDP-glucuronosyltransferases) enzymes, guanylate cyclases, glyoxylate and dicarboxylate metabolism, and amino acid metabolism, such as arginine and proline metabolism and glutamate/ glutamine metabolism (**Appendix Fig S1**). Examples of Q4 genes that are highly variant both during development and in different tissues include *ugt-13, ugt-18* and *ugt-34* (UGT enzymes); *gcy-3* (guanylate cyclases); and *ddo-3, gln-2, argk-1* and *phy-3* (amino acid metabolism) (**Fig 1I**).

Altogether, our analyses indicate that at least 75% (1,867 out of 2,492) of metabolic genes vary in mRNA levels and are therefore likely transcriptionally regulated, including 902 iCEL1314 genes (∼69%) (**Fig 1B**). These numbers are similar to the proportion of varying non-metabolic genes (∼68%), indicating that metabolic genes are overall at least as much under transcriptional control as other genes (**Fig 1B**). Interestingly, there are differences among different types of metabolic genes. For instance, amino acid metabolism genes are variant in both development and in tissues, while lipid metabolism genes are mostly variant in tissues, and growth and energy metabolism are relatively invariant in both development and in tissues.

### A Supervised Approach Shows Widespread Coexpression of Genes Comprising Metabolic Pathways

Since most metabolic genes exhibit variation in gene expression and because genes that are coexpressed often function together (Eisen et al., 1998; Hughes et al., 2000; Kim et al., 2001; Segal et al., 2003; Stuart et al., 2003), we investigated whether genes that function in the same metabolic pathway tend to be coexpressed. We previously found that the five genes comprising the propionate shunt pathway are coordinately activated in response to propionate accumulation (Bulcha et al., 2019; Watson et al., 2016). In addition, we found strong coexpression of genes functioning in the methionine/S-adenosylmethionine (Met/SAM) cycle, for instance when flux through this pathway is perturbed (Giese et al., 2020). To systematically test which *C. elegans* metabolic pathways exhibit coexpression, we developed a custom pathway enrichment analysis pipeline (**Fig 2A**) based on gene set enrichment analysis (GSEA) (see **Methods**) (Subramanian et al., 2005) and applied it to the compendium. We used mean coexpression of all metabolic genes with the genes in a given set (*i.e.,* a pathway) as our ranking metric and calculated an enrichment score (ES) that defines the enrichment of relatively high coexpression within that set. A normalized ES (NES) indicates relative strength of this enrichment compared to randomized tests, the significance of which is measured as a false discovery rate (FDR) (**Fig 2A**).

**Figure 2.**
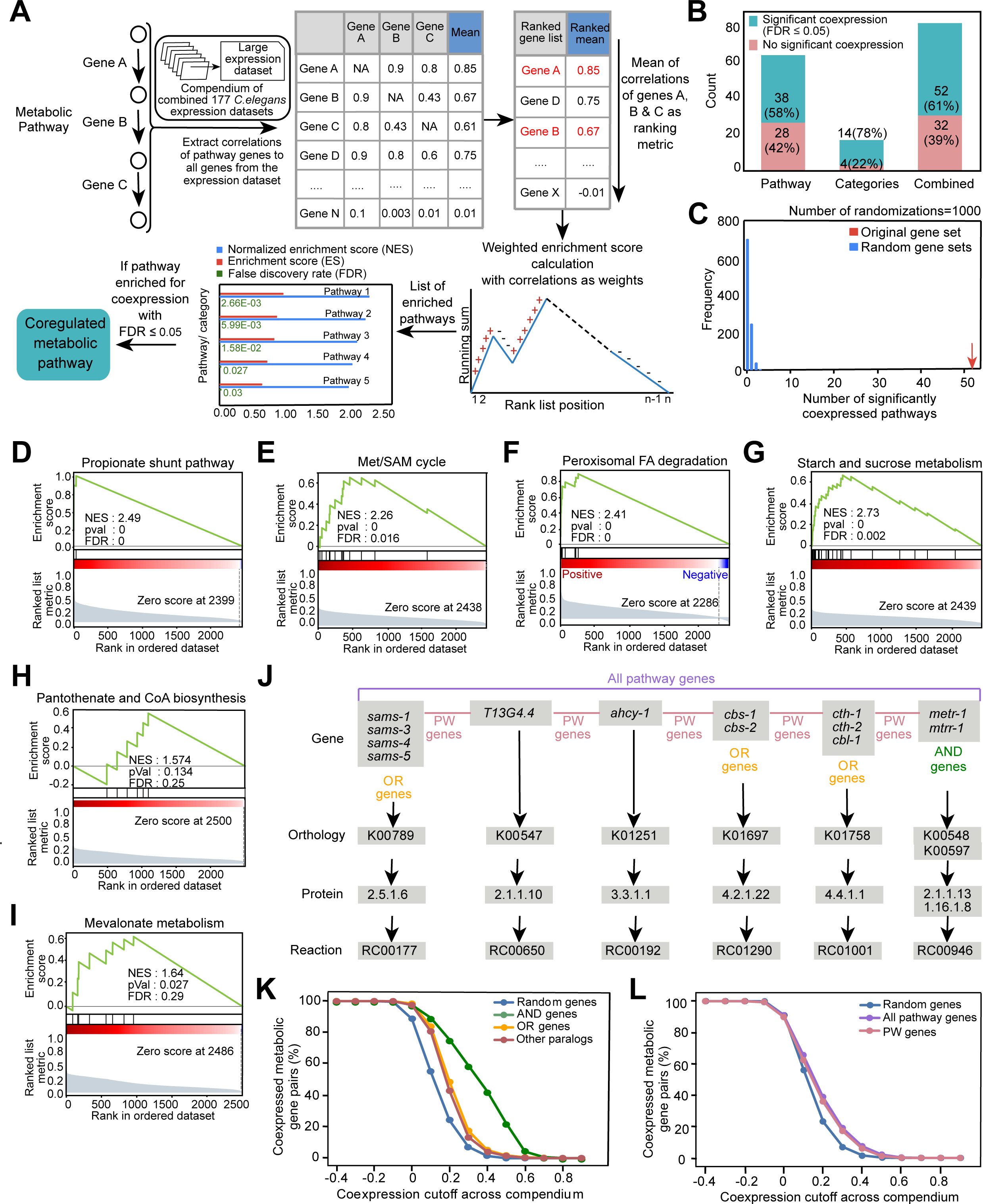
Supervised Approach to Investigate Coexpression of Metabolic Pathways. (A) Custom computational pathway enrichment analysis pipeline that determines coexpressed genes functioning in the same metabolic pathway. Pairwise coexpression was based on the gene expression compendium. For every annotated metabolic pathway, the coexpression of pathway genes (columns) to all metabolic genes (rows) was extracted. A ranked list of genes was obtained for each pathway by taking the mean of coexpression values in rows while ignoring self-correlations. Weighted gene-set enrichment analysis was then performed to find significantly enriched pathways. If a pathway is self-enriched with FDR ≤ 0.05, it is annotated as coexpressed. (B) Bar graph indicating the percentage of metabolic pathways and categories that show significant coexpression compared to ones that are not self-enriched for coexpression. (C) Histogram denoting the number of significantly coexpressed metabolic pathways obtained by 1000 randomizations while maintaining the structure of the data. (D-I) Mountain plots showing self-enrichment of (D) propionate shunt pathway, (E) Met/SAM cycle, (F) peroxisomal fatty acid degradation pathway, (G) starch and sucrose metabolism, (H) pantothenate and CoA biosynthesis, and (I) mevalonate metabolism (J) Metabolic pathways often consist of reactions catalyzed by single genes, OR genes and AND genes. All genes involved in the same pathway are collectively annotated as all pathway genes. Genes that are associated with distinct reactions are annotated as PW genes. PW gene pairs exclude AND and OR gene pairs. Met/SAM cycle pathway, which consists of 13 metabolic genes, is shown as an example. (K) Percentages of pairs of AND genes, OR genes, other paralogs, and random metabolic genes categorized as coexpressed using different coexpression values as cutoffs. Coexpression values are based on the gene expression compendium. (L) Percentage of random, all pathway, and PW gene pairs categorized as coexpressed using different coexpression values as cutoffs. Coexpression values are based on the gene expression compendium.

We used this tool to investigate the prevalence of coexpression in annotated metabolic pathways, enzyme complexes, and enzyme families as defined in WormPaths (Walker et al., 2021). Henceforth, we use ‘category’ to refer to a group of metabolic genes that best fit in an enzyme complex or related set of enzymes such as amino-acyl-tRNA synthetases, rather than a true pathway in which metabolites are sequentially converted (Walker et al., 2021). With an FDR cutoff of ≤ 0.05, 52 of 84 metabolic pathways or categories (∼61%) exhibit coexpression (**Fig 2B, Table EV7, Appendix Fig S2**). We validated this result by running the custom pathway enrichment analysis pipeline on 1,000 randomized gene sets. For these randomizations, the structure of the data was maintained such that the correlation matrix, the number of genes in each pathway and the number of times an individual gene was repeated across pathways all remained the same. The number of pathways or categories found to be enriched for coexpression with 1,000 randomizations was consistently very small in randomized data compared to the number from the original gene sets (**Fig 2C**). This validates the observation that genes in a metabolic pathway tend to be coexpressed. As expected, the 52 coexpressed metabolic pathways and categories include the propionate shunt and the Met/SAM cycle (**Fig 2D-E**). When we examined coexpression in pathways and categories separately we found that 78% of categories showed significant coexpression compared to 58% of pathways (**Fig 2B**). Examples of metabolic pathways that exhibit high coexpression include peroxisomal FA degradation and starch and sucrose metabolism (**Figs 2F and 2G**). Examples of coexpressed categories include vacuolar ATPases, ETC complex I, and aminoacyl-tRNA synthetases (**Appendix Fig S2**). There are 32 categories and pathways that do not exhibit self-enrichment, including pantothenate and CoA biosynthesis and mevalonate metabolism (**Figs 2H and 2I, Appendix Fig S2**). Such pathways may either not be regulated at all, may be regulated by allostery, or only one or a few genes in these pathways are transcriptionally regulated and may therefore function as key regulatory genes.

The extent of within-pathway coexpression of metabolic genes can be potentially confounded because metabolic reactions are often associated with multiple genes in gene-protein-reaction (GPR) annotations (Kim et al., 2008; Thiele and Palsson, 2010). There are two reasons for this. First, some metabolic reactions are catalyzed by enzyme complexes comprising two or more proteins. In such cases, all genes need to be expressed for the reaction to take place and are therefore annotated here as ‘AND’ genes. Second, some genes are part of larger families (paralogs) that encode isozymes or highly similar proteins. Metabolic network reconstruction efforts use protein sequence homology to associate genes with reactions (Thiele and Palsson, 2010) (Yilmaz and Walhout, 2016, 2017). As a result, multiple highly homologous paralogs may be associated with the same metabolic reaction. Such paralogs are annotated here as ‘OR’ genes. Some reactions are associated with a combination of AND and OR genes (**Fig EV4A**). Finally, for some gene families it may be that one member catalyzes one reaction and another member catalyzes another. Paralogs that are associated with distinct reactions are referred to here as “other paralogs” (**Fig EV4B**). Pathways can be associated with multiple types of AND and OR genes (**Fig 2J**).

AND genes encode proteins that function together in complexes, and it is well-known that such genes are often strongly co-expressed (Jansen et al., 2002). For example, genes encoding ETC complex members are coexpressed and coregulated (van Waveren and Moraes, 2008). Therefore, we wondered if this holds true for AND genes in iCEL1314 and, if so, whether this would inflate pathway coexpression enrichment. To test this, we systematically assessed coexpression of different types of gene pairs. As expected, we found that AND genes are significantly more coexpressed than random gene pairs, OR genes and other paralogs (**Figs 2K, EV4C, EV4D, EV4E, EV4F and EV4G**). Both OR genes and other paralogs are also more coexpressed than random metabolic gene pairs in all three datasets (**Figs 2K, EV4C, EV4D, EV4E, EV4F and EV4G**). Surprisingly, OR genes are more coexpressed than other paralogs across tissues (**Figs 2K, EV4C, EV4D, EV4E, EV4F and EV4G**).

Based on the analysis of AND and OR genes, it is difficult to determine the contribution of the coexpression of such gene pairs to pathway enrichment. Therefore, we examined coexpression of gene pairs that are annotated with distinct reactions in a pathway, which we refer to as pathway (PW) genes (**Fig 2J**). We found that PW gene pairs are also significantly more coexpressed than random gene pairs (**Figs 2L, EV4H, EV4I, EV4J and EV4K**). Therefore, pathway coexpression is not just driven by OR and AND genes, indicating that enzyme coexpression is a feature of many metabolic pathways.

### An Unsupervised Approach Extracts Coexpressed Sub-Pathways

Our finding that metabolic pathways and categories exhibit extensive coexpression was based on previously annotated pathways (Walker et al., 2021). However, these pathways connect into the larger metabolic network and the definition of the start and ending of each pathway is somewhat arbitrary. Since there is extensive coexpression of genes that function together in pre-defined pathways, we reasoned that we may be able to use coexpression to extract functionally connected metabolic genes or pathways in an unbiased manner.

Reactions in linear pathways have complete flux dependence, or ‘coflux’ (*i.e.*, coflux = 1 for all pairs of reactions in a linear pathway), while in branched pathways flux dependency may be partial (coflux = between 0 and 1), and in uncoupled reactions there is no dependence (coflux=0). It has been previously shown in *E. coli* that flux coupling is a better indicator of coregulation among the corresponding metabolic genes than their absolute distance in the metabolic network (Notebaart et al., 2008). Therefore, we developed a custom unsupervised approach to identify sets of metabolic genes that are annotated to flux-dependent reactions in iCEL1314 and that are highly coexpressed in the gene expression compendium (**Fig 3A, Tables EV8, EV9 and EV10**) (details in **Methods**). The coflux algorithm uses FBA to simulate reaction fluxes and measure flux dependency between reaction pairs and uses GPR rules to convert the resulting reaction coflux matrix to a gene coflux matrix (see **Methods**). To define metabolic genes that are highly coexpressed, we used the coexpression matrix obtained from the gene expression compendium. We then multiplied the coflux and coexpression matrices to obtain a ‘product matrix’, which has large product values only for gene pairs that are both coexpressed and share flux dependency (**Fig 3B**).

**Figure 3.**
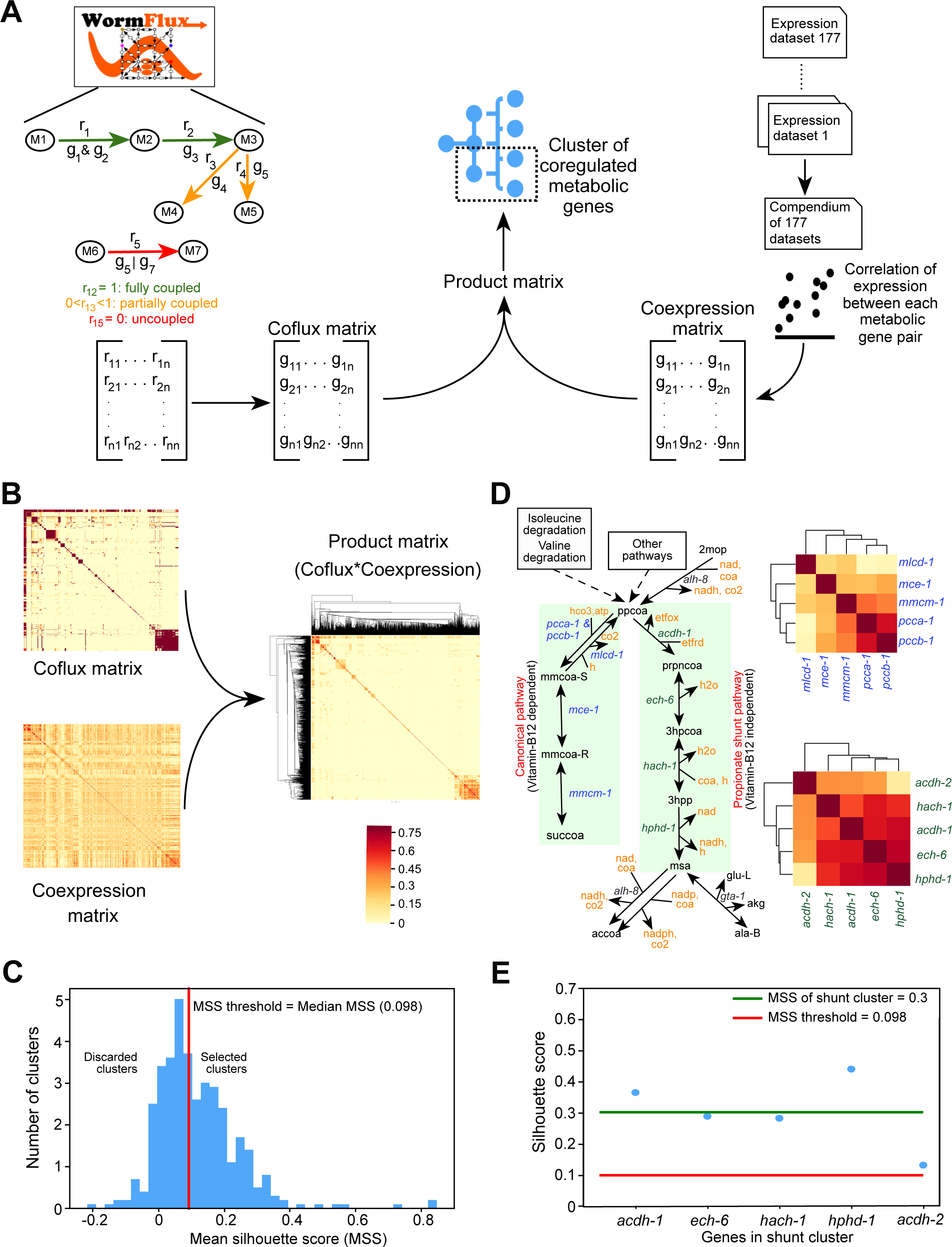
An Unsupervised Approach to Extract Coexpressed and Flux-Dependent Metabolic Genes. (A) Computational pipeline to extract tightly coregulated units in the metabolic network: Functional relationships are provided through theoretical flux associations (coflux) calculated using *C. elegans* metabolic network model iCEL1314, while expression correlations come from the compendium of 177 expression datasets. Hierarchical clustering on the product of coflux and coexpression matrix gives coexpressed metabolic pathways. (B) Heatmaps showing coflux and coexpression of iCEL1314 genes and clustered heatmap showing added modularity to coexpression space by product of coexpression and coflux. (C) Plot showing the distribution of the MSS of all clusters obtained using dynamic cut tree algorithm with stringent parameters (deepSplit=2, minClusterSize=3). Median of MSS of all clusters was chosen as a threshold to define high-quality clusters for analysis. (D) Distinct clusters denoted by clustered heatmap of genes in canonical and shunt pathways of propionate degradation were extracted using dynamic cut tree algorithm with stringent parameters (deepSplit=2, minClusterSize=3). (E) Scatter plot with individual silhouette scores of genes in the propionate shunt cluster. MSS of this pathway is shown along with the overall MSS threshold.

To extract coexpressed sub-pathways, we clustered the product matrix using the dynamic cut tree algorithm (Langfelder et al., 2008) with a relatively stringent set of parameters (see **Methods** for details). This led to 197 clusters, ranging in size from three to 27 genes. To evaluate cluster quality, we determined the cohesion of genes within a cluster and the separateness from genes in other clusters using a Silhouette Score (SS) (Rousseeuw, 1987). We then used the mean SS (MSS) of each cluster and used the median of the MSS distribution (MSS=0.098) as a threshold to extract 99 high-quality clusters (**Fig 3C, Table EV9**).

The propionate shunt pathway genes formed a tight cluster (**Figs 3D and 3E**). Interestingly, while the first four genes, *acdh-1, ech-6, hach-1* and *hphd-1,* occurred closely together, the fifth gene, *alh-8*, was not part of the same cluster. This could be explained in two ways. First, *alh-8* encodes an enzyme that functions at a junction in the pathway where its substrate malonate semialdehyde is converted either to acetyl-coa or, potentially, to beta-alanine. Therefore, metabolic flux is divided in two directions, and is not linearly coupled with the shunt pathway flux like the first four reactions. Second, *alh-8* is annotated to another reaction where 2-methyl-3-oxopropionate is converted to propionyl-CoA (**Fig 3D**). This clustering approach also revealed coexpression of genes comprising the canonical, vitamin B12-dependent propionate degradation pathway, indicating that this pathway may be transcriptionally activated or repressed under specific conditions (**Fig 3D, Table EV9).**

### The Unsupervised Approach Reveals Pathway Boundaries

As stated above, all pathways are connected to form the metabolic network and pathway boundary definitions are somewhat arbitrary. The propionate shunt example above shows that the unsupervised approach can extract sub-pathways (*e.g.*, the propionate shunt) from arbitrarily defined pathways (*e.g.*, propionate degradation) based on coexpression and flux dependency. We therefore used other clusters defined by the unsupervised approach to better define functional starts and ends of different pathways.

An example of a pre-annotated WormPaths pathway that was fully captured with the unsupervised approach is peroxisomal FA degradation, where genes from each reaction formed a high-quality cluster (MSS=0.23) (**Fig 4A, Table EV9**). Therefore, in this case, the pathway boundaries cover the entire original pathway. The first reaction in peroxisomal beta-oxidation is catalyzed by acyl-CoA oxidases (encoded by *acox* genes). Only *acox-1.1* and *acox-3* in the *acox*-family genes are coexpressed with the other peroxisomal FA oxidation genes, indicating that they are more likely to function in this pathway than the other *acox* genes, which are coexpressed with each other, and with mitochondrial FA degradation genes (**Table EV9**).

**Figure 4.**
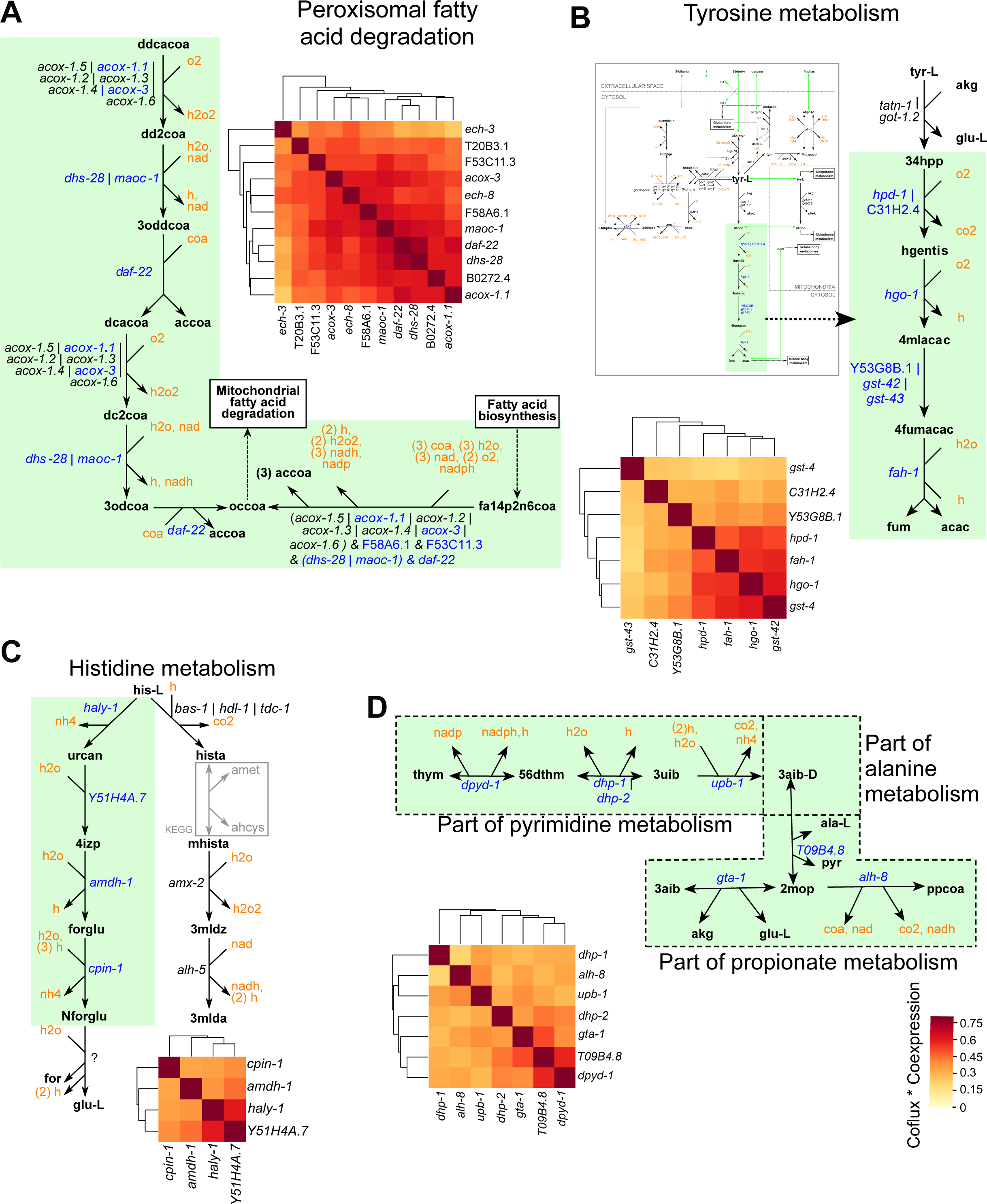
An Unsupervised Approach Defines Metabolic Pathway Boundaries. (A) Clustered heatmap denoting a distinct cluster consisting of at least one gene from every reaction in peroxisomal fatty acid degradation. Heatmap genes are shown in bold font. (B) Clustered heatmap showing a distinct cluster formed by the tyrosine degradation genes separate from the rest of the tyrosine metabolism. (C) Clustered heatmap showing a distinct cluster formed by a boundary within the histidine degradation pathway. (D) Clustered heatmap showing a distinct cluster formed by genes traversing pathway boundaries that are parts of propionate, alanine and pyrimidine metabolism.

Tyrosine metabolism offers an example where only a subset of annotated genes clustered together. Tyrosine can be metabolized via different reactions, in different pathway branches (**Fig 4B**). In one pathway branch, tyrosine is degraded in five steps to produce fumarate and acetoacetate. The genes in this branch; *gst-43, C31H2.4, Y53G8B.1, hpd-1, hgo-1, fah-1 & gst-42* form a tight cluster with MSS=0.358 (**Fig 4B, Table EV9**). Some of these genes are OR genes such *as hpd-1* OR C31H2.4; and Y53G8B.1 OR *gst-42* OR *gst-43*, which suggests that these genes are correctly annotated to this pathway branch.

Histidine can also be degraded via two pathway branches: one converting histidine to glutamate through N-formyl-L-glutamate, and the other converting histidine to 3-methylimidazoleacetic acid. The four genes associated with the conversion of histidine to N-formyl-L-glutamate; *haly-1, Y51H4A.7, amdh-1 and cpin-1,* form one of the top-ranked clusters (MSS = 0.357) (**Fig 4C**). However, the genes in the other branch are not coexpressed. Therefore, the unsupervised clustering method identified coexpressed branches, or sub-pathways, of both tyrosine and histidine degradation, illustrating its utility as an addition to the predefined metabolic pathways in WormPaths.

We found that *alh-8* and *gta-1*, which are functionally associated but not coexpressed with the propionate shunt (**Fig 3D**), cluster with T09B4.8 (alanine metabolism) and four other genes belonging to pyrimidine metabolism: *dpyd-1, dhp-1, dhp-2* and *upb-1* (**Fig 4D, Table EV9**). This observation functionally connects genes in what were heretofore separately annotated pathways, *i.e.*, pyrimidine, alanine, and propionate metabolism. The coexpression of these genes suggests that thymine is degraded, leading to the formation of propionyl-CoA, L-3-amino-isobutanoate and acetyl-CoA. This observation also suggests that *alh-8* levels have a stronger functional role in the conversion of 2-methyloxopropanoate to propionyl-CoA than in the propionate shunt. Remarkably, this further indicates that *alh-8* may participate in both the generation and degradation of propionate.

The stringent parameters we used in the dynamic tree cut algorithm favor small clusters, and as a result, interconnections between different pathways may be lost in the analysis. To unveil such connections, we relaxed the parameters to allow the derivation of larger clusters of coexpressed genes (**Table S10, Fig EV5A**). With these settings, the propionate shunt cluster expanded and included *bckd-1A, bckd-1B, dbt-1, Y43F4A.4, ard-1, acdh-3, acdh-9* and *B0250.5,* which are annotated to branched chain amino acids (BCAA) isoleucine and valine degradation pathways, but not *alh-8* or *gta-1* (**Fig EV5B, Table S10**). Propionyl-CoA, the starting metabolite of the propionate shunt, is produced by the breakdown of valine and isoleucine. We recently proposed that the propionate shunt not only functions to detoxify excess propionate, but also to produce acetyl-CoA for ketone body and energy production (Ponomarova et al., 2022). The coexpression of valine and isoleucine breakdown genes with the propionate shunt indicates a functional connection between these pathways to produce energy.

The Met/SAM cycle provides another example of different degrees of clustering that can be unveiled with different parameter settings (**Fig EV5C**). With relaxed settings, many pathway genes were annotated to the same cluster, while with stringent settings four smaller clusters were defined. The smaller clusters captured different parts of one-carbon metabolism with their connections to Met/SAM cycle. One of the clusters consisted of *adk-1, pmt-1*, and *pmt-2*, along with the Met/SAM cycle genes *ahcy-1* and *sams-1*. This coexpression connects the Met/SAM cycle with glycerophospholipid metabolism, specifically phosphatidylcholine biosynthesis, which depends on methylation reactions using SAM (*pmt-1* and *pmt-2*)(Walker et al., 2011), as well as with purine metabolism (*adk-1*) (Ducker and Rabinowitz, 2017). A second stringent cluster shows that Met/SAM cycle genes are highly coexpressed with the folate cycle gene *mthf-1*. MTHF-1 produces the methyl group that is used to convert homocysteine into methionine in the Met/SAM cycle (Ducker and Rabinowitz, 2017; Giese et al., 2020). The folate-cycle related genes are also highly coexpressed with *chdh-1*, which encodes a dehydrogenase that converts choline to betaine aldehyde. Together, these results confirm that the Met/SAM cycle is overall coexpressed (Giese et al., 2020) and show that additional co-functioning genes can be identified.

The unsupervised approach also identified gene clusters that are not part of any coexpressed pathway identified by the supervised method above. An example is selenocompound metabolism, where a set of seven genes; *mett-10, pstk-1, trxr-1, pps-1, secs-1, nsun-5* and *seld-1* form a highly coexpressed cluster (FDR=0.025, NES=1.7) (**Fig EV5D**). In comparison, the respective FDR and NES values for self-enrichment of the WormPaths pathway of selenocompound metabolism were 0.75 and 0.98 (**Table EV7**). Altogether, these results illustrate that not all genes in a pathway are coexpressed and further indicate that a subset of a pathway or a combination of subsets from multiple pathways may be under transcriptional control.

### Metabolic Pathway Communities Reveal Coexpression Among Complexes and Pathways

To explore additional coexpression clusters than those that were captured by the relaxed settings described above, we visually inspected the product matrix and extracted four clusters we refer to as metabolic pathway ‘communities’ (**Fig 5A**). We analyzed these communities by WormPaths PEA (Walker et al., 2021). The first community is enriched in ETC complexes I, III and IV, indicating broad transcriptional control of energy production (**Fig 5B, Table EV11**). The second community is enriched in mitochondrial and peroxisomal FA degradation, FA biosynthesis, ascaroside biosynthesis, and BCAA degradation (**Fig 5C, Table EV11**). The connection between FA metabolism and BCAA degradation may reflect the fact that some FAs are synthesized from BCAA breakdown products. For instance, branched-chain fatty acids (BCFAs) are synthesized from the branched-chain alpha keto acids of valine, leucine and isoleucine such as isovaleryl-CoA and isobutyryl-CoA after their decarboxylation and further chain elongation (Daschner et al., 2001; Jia et al., 2016; Wallace et al., 2018). The third community is enriched in UGT enzymes and pentose and glucuronate interconversions (**Fig 5D, Table EV11**). UGT enzymes are known to conjugate glucuronic acid to facilitate the elimination of toxic substances from the body (King et al., 2000). This result suggests that this type of detoxification in *C. elegans* is transcriptionally coordinated. The fourth community is enriched in aminoacyl-tRNA biosynthesis, N-glycan biosynthesis, collagen biosynthesis, iron metabolism, and mevalonate metabolism, all of which produce biomass precursors. While aminoacyl-tRNA synthetases play a major role in protein biosynthesis by linking amino acids to their cognate transfer RNAs (tRNAs), mevalonate metabolism provides precursors for glycan, collagen biosynthesis provides collagen for formation of the cuticle and other extracellular matrices, and iron metabolism is important for many aspects of metabolism, including the production of heme groups of heme proteins. This result points toward the possibility that growth is transcriptionally regulated by a central mechanism controlling pathways that produce biomass precursors and assemble biomass (**Fig 5E, Table EV11**). Taken together, we confirmed the coexpression of metabolic pathways, revealed coexpressed sub-pathways, as well as coexpression among pathways.

**Figure 5.**
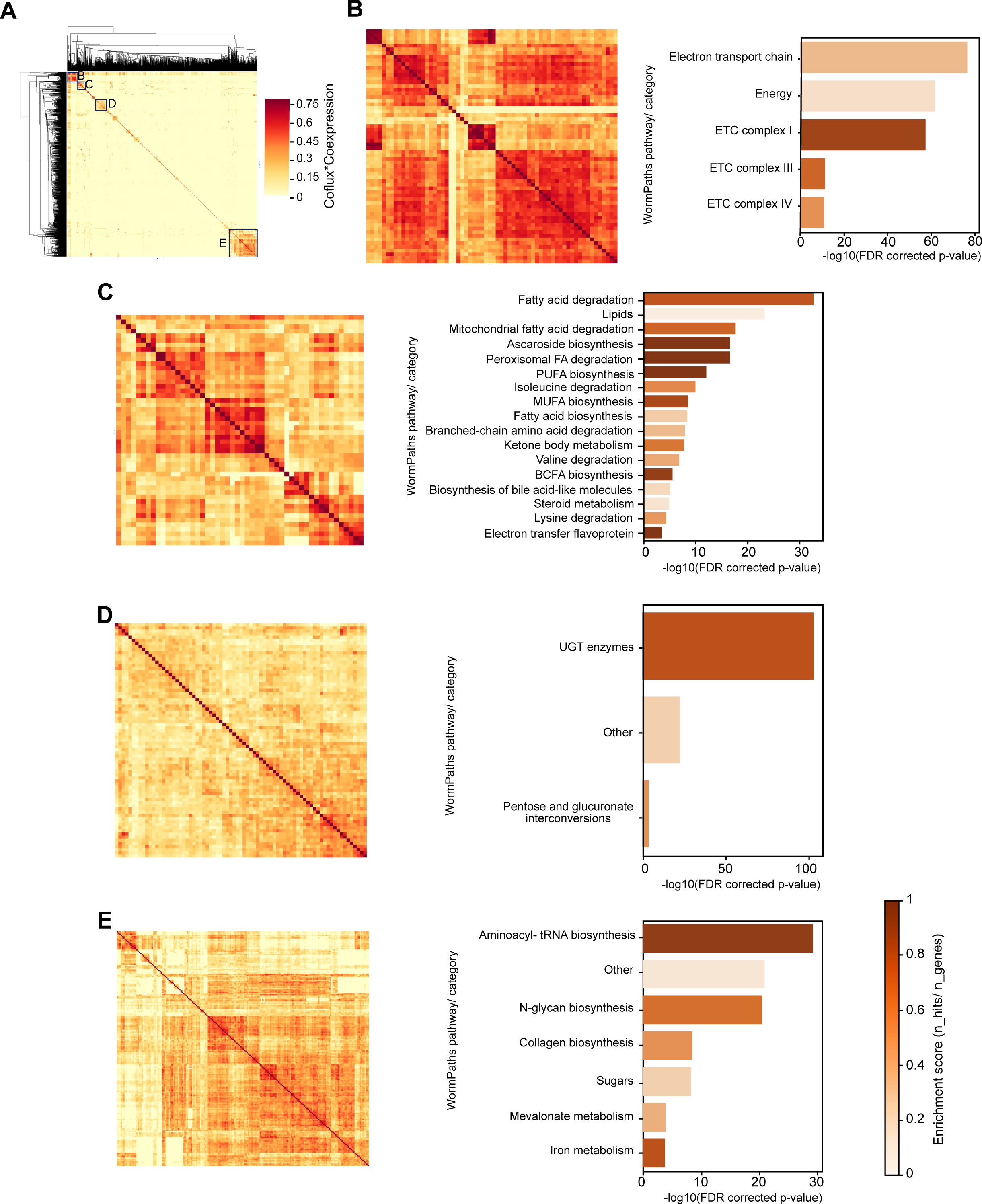
Extraction of Metabolic Communities. (A) Clustered heatmap indicating communities formed by multiplying coflux and coexpression values of iCEL1314 genes. B, C, D and E define four major communities shown in the respective parts of this figure. (B-E) PEA of communities B (B), C (C), D (D) and E (E).

### Metabolic Sub-Pathways are Activated or Repressed under Different Conditions

The gene expression compendium is comprised of 177 expression profiling datasets that measure relative mRNA levels in a variety of experimental conditions and genotypes. Therefore, we next asked whether we could identify specific conditions in which different metabolic gene clusters are activated or repressed. Using a custom computational pipeline (**Fig 6A**), we first identified those datasets that best represent the coexpression of a particular cluster. We then manually investigated the top datasets for each cluster (see **Methods**). To validate this approach, we first examined the expression of propionate shunt cluster genes in top-scoring datasets. We previously showed that propionate shunt genes are repressed in animals fed *Comamonas aquatica* DA1877, a bacterium that (unlike the standard *E. coli* OP50 diet) produces vitamin B12 thus enabling flux through the canonical propionate degradation pathway (MacNeil et al., 2013; Watson et al., 2013; Watson et al., 2014; Watson et al., 2016). The dataset from that study, labeled as dataset 15 in the compendium, scored as most significant for propionate shunt gene coexpression, where the genes are expressed in animals fed *E. coli* OP50, but not in animals fed *C. aquatica* (**Fig 6B, Table EV4**). We found a similar trend in dataset 51 (Miller et al., 2015), where propionate shunt genes are repressed in animals fed *Pseudomonas aeruginosa*, which also provides vitamin B12 (Watson et al., 2014) (**Fig 6B**). Interestingly, we also found that propionate shunt genes are highly coexpressed in a dataset that measured expression in *spr-5* mutants versus wild type animals across 1(f1), 13(f13) and 26(f26) generations (**Fig 6B**, dataset 139). Propionate shunt genes are more highly expressed in the N2 reference strain compared to *spr-5* mutant animals (**Fig 6B**). Since *spr-5* encodes a histone demethylase, this may indicate that the expression of shunt genes is regulated not only by TFs, but also by epigenetic mechanisms.

**Figure 6.**
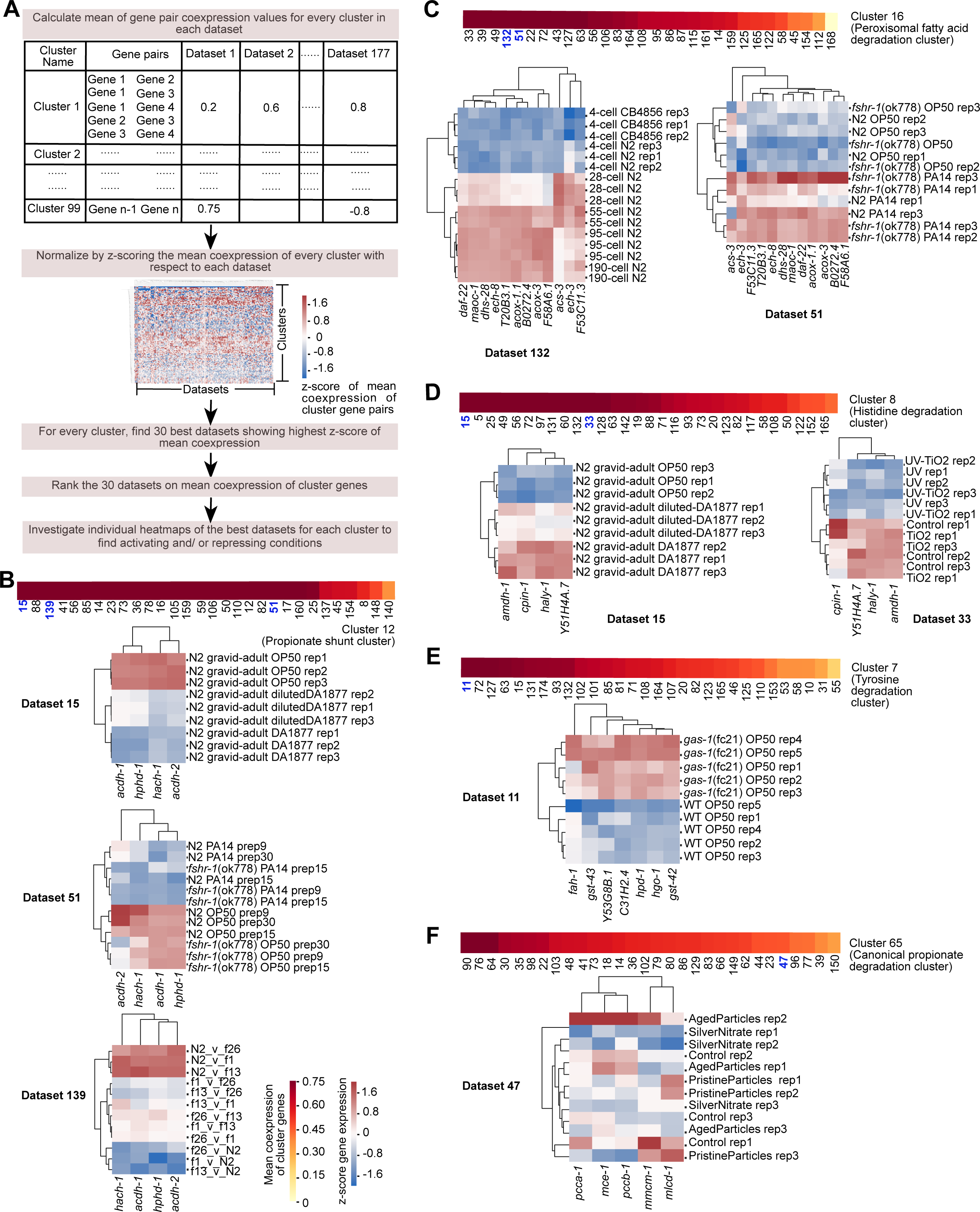
Condition Analysis of Metabolic Gene Coexpression. (A) Computational pipeline to extract activating or repressing conditions of metabolic clusters. The mean coexpression of all gene pairs in each cluster in each dataset are calculated separately. To rank datasets that showed highest coexpression uniquely for each cluster, these mean coexpression values are normalized using z-scoring across each dataset (as shown by heatmap). 30 best datasets that potentially represent activation/repression conditions of each cluster are identified by the z-score values of mean coexpression. Then, for each cluster, datasets are manually inspected in the order of decreasing mean coexpression along with its associated published paper and the heatmap to understand activation/repression conditions. (B-F) Mean coexpression of the 30 best datasets for clusters of propionate shunt (B), peroxisomal fatty acid degradation (C), histidine degradation (D), tyrosine degradation (E), and canonical propionate degradation (F), followed by heatmap examples from selected datasets as indicated by bold-blue dataset numbers. Color bar and heat map legend as indicated in (B).

Peroxisomal FA degradation genes were most significantly coexpressed in dataset 132, which measured gene expression in precisely staged embryos during the first quarter of embryonic development (**Fig 6C**). The time course included a stage of exclusively maternal transcripts (four-cell), the transition to zygotic transcription (28-cell), and the presumptive commitment to the major cell fates (55-, 95-, and 190-cell stages) (Yanai and Hunter, 2009). Peroxisomal FA degradation genes were lowly expressed in four-cell embryos and their expression increased in later embryonic stages, which may reflect a change in carbon source for energy and biomass generation prior to hatching and feeding. Peroxisomal FA degradation genes are also upregulated in animals fed *P. aeruginosa* compared to the standard *E. coli* OP50 diet (**Fig 6C**, dataset 51). This may reflect the high energy demand during infection (Nhan et al., 2019).

Inspection of expression of the histidine degradation cluster discussed above revealed that it is highly expressed in animals fed *C. aquatica* (**Fig 6D**, dataset 15). However, these genes were not affected in animals fed *E. coli* supplemented with vitamin B12 (Bulcha et al., 2019), suggesting that the effect of *C. aquatica* on this cluster may be independent of this cofactor. Animals showed lower expression of this cluster upon exposure to UV treatment (**Fig 6D**, dataset 33). In humans, UV converts cis-uruconate (the product of first reaction of histidine degradation, **Fig 4C**) to trans-uruconate, which has been proposed to play a protective role in skin (Brosnan and Brosnan, 2020). Our result indicates that, in *C. elegans*, UV exposure rewires metabolic flux to avoid histidine degradation, for instance to preserve histidine that could be converted to trans-uruconate.

The tyrosine degradation cluster was most significantly coexpressed in dataset 11, wherein expression profiles of *gas-1* mutant animals, which are deficient in mitochondrial respiration, were compared to wild type animals (Falk et al., 2008) (**Fig 6E**). This study revealed that free tyrosine levels are decreased in *gas-1* mutants. In addition, there is a failure of NAD+-dependent ketoacid oxidation in mitochondrial respiratory chain mutants (Falk et al., 2008). To compensate for this respiratory dysfunction, multiple pathways are upregulated, including the TCA cycle and ketone body metabolism (Falk et al., 2008). Since the end products of tyrosine degradation pathway cluster are the TCA cycle intermediate fumarate and the ketone body acetoacetate (**Fig 4B**), the function of the upregulation of tyrosine degradation during mitochondrial dysfunction may be to supply metabolites for compensatory pathways.

These results show that the gene expression compendium can be used to gain insight into the conditions that most greatly affect the activation or repression of different metabolic gene clusters. However, one needs to be careful to manually inspect the conditions of interest because sometimes coexpression can be biased by an outlier experiment. An example of this is dataset 47, one of the top datasets in which canonical propionate degradation pathway genes are coexpressed. Even though the conditions in this dataset are not related to canonical propionate breakdown, this dataset falsely appears as one of the top datasets due to coexpression driven by one bad outlier sample. (**Fig 6F**).

Overall, our systematic analysis revealed specific conditions of when metabolic gene clusters are activated or repressed, reinforcing our overall finding that transcriptional regulation plays an important role in the control of metabolism.

### WormClust Web Application Enables Gene-By-Gene Query To Identify Coexpression with Metabolic (Sub)-Pathways

A major premise of this study is the assumption that variance in mRNA levels results, at least in part, from transcriptional regulation, which in turn suggests that genes are coexpressed because they are coregulated. In reverse engineering of gene regulatory networks, coexpression of TFs with their target genes has been used to define causal relationships (MacNeil and Walhout, 2011; Segal et al., 2004). To make our data available to the community as well as to enable easy searching of coexpression of TFs and other non-metabolic genes with metabolic (sub)-pathways, we developed a web-application we named WormClust, which is available on the WormFlux website (Yilmaz and Walhout, 2016). This tool takes any *C. elegans* gene as input and evaluates its coexpression with metabolic (sub)-pathways. If the query gene is an iCEL1314 gene, the output is a clustered heatmap of the coexpressed genes in the model based on product matrix, and according to the selected level of stringency, i.e., relaxed or stringent. If the query gene is not an iCEL1314 gene, then an association of the gene with annotated metabolic pathways is provided. The threshold for this association can be based on FDR and/or NES (**Fig 7A**).

**Figure 7.**
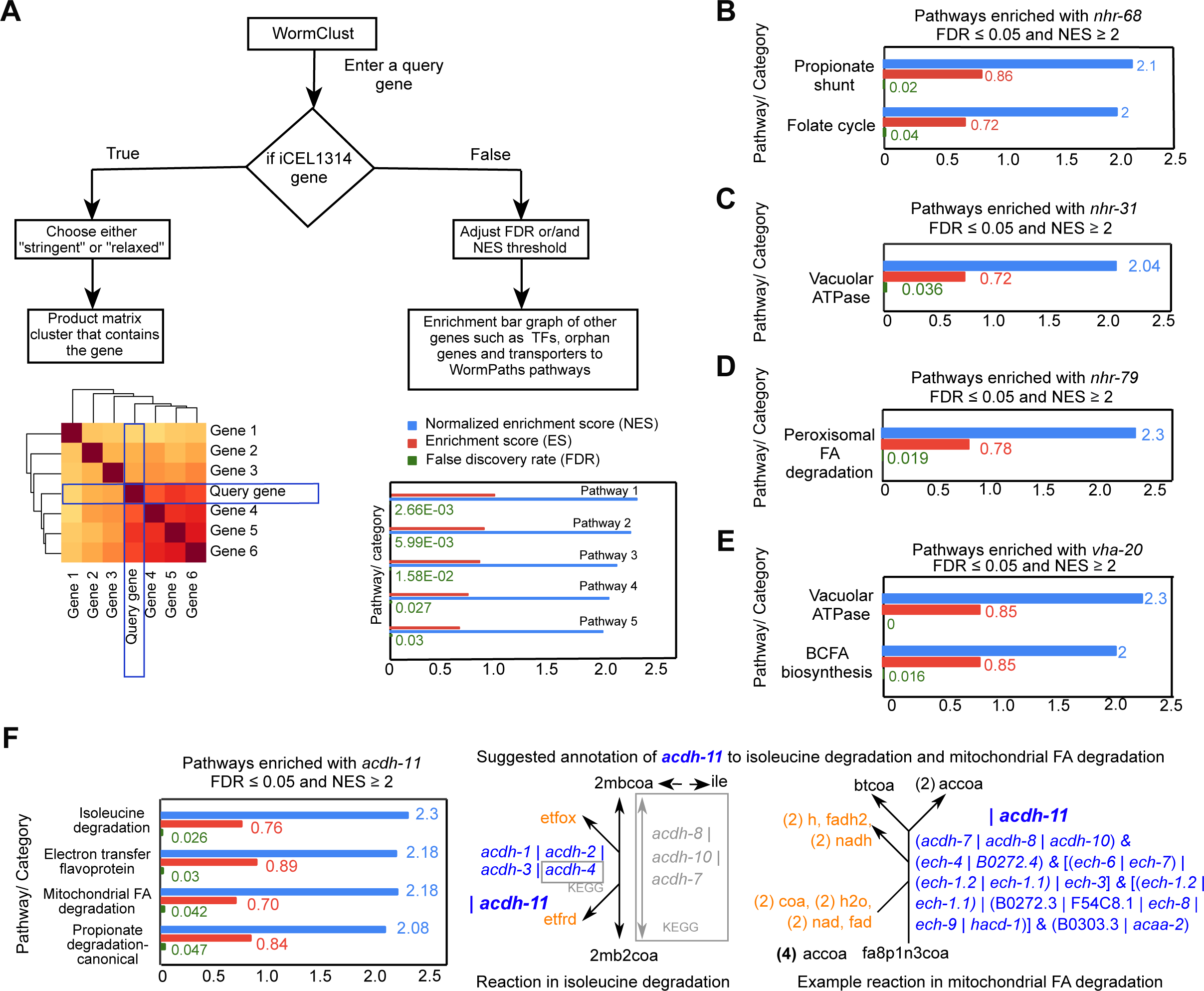
WormClust: A Web Application that Enables Querying of Genes To Identify Coexpression with Metabolic (Sub)-Pathways. (A) Diagram showing workflow of WormClust. A *C. elegans* gene is taken as input. If the gene is part of iCEL1314, a clustered heatmap of closely associated genes in product matrix of coflux and coexpression is displayed, based on stringency level of clustering. If the input gene is not an iCEL1314 gene, enrichment bar-graphs of the gene to annotated metabolic pathways are displayed, based on selected FDR and NES thresholds. (B) Bar graph of pathways that are significantly coexpressed with *nhr-68* with NES ≥ 2 and FDR ≤ 0.05. (C) Plot showing significant coexpression of *nhr-31* with vacuolar ATPases (FDR≤ 0.05 and NES ≥ 2). (D) Plot showing significant coexpression of *nhr-79* with peroxisomal fatty acid degradation (FDR≤ 0.05 and NES ≥ 2). (E) Plot showing significant coexpression of *vha-20* with vacuolar ATPases (FDR≤ 0.05 and NES ≥ 2). (F) Bar graph of pathways that are significantly coexpressed with *acdh-11* with NES ≥ 2 and FDR ≤ 0.05 (left). Examples of specific reactions in metabolic network model where *acdh-11* can be annotated as OR gene (right).

We, and others, previously found that nuclear hormone receptor (NHRs) are TFs that frequently associate with metabolic genes in different types of assays and dataset (Arda et al., 2010; Bhattacharya et al., 2022; Mori et al., 2017; Van Gilst et al., 2005). To illustrate the utility of WormClust, we tested three NHRs with known metabolic pathway associations for coexpression with annotated metabolic pathways in the compendium of 177 *C. elegans* expression datasets. All of these showed coexpression with their target metabolic pathways (**Figs 7B, 7C and 7D**): *nhr-68* was highly coexpressed with the propionate shunt (FDR=0.02 and NES=2.1) (Bulcha et al., 2019), *nhr-31* associated with vacuolar ATPases (FDR=0.036, NES=2.04) (Hahn-Windgassen and Van Gilst, 2009), and *nhr-79* is coexpressed with peroxisomal FA degradation (Zeng et al., 2021).

WormClust also provides an opportunity to annotate new metabolic genes. For example, *vha-20,* a vacuolar ATPase that is not part of the original iCEL model because it was only recently annotated by WormBase and KEGG, shows significant enrichment to vacuolar ATPases (**Fig 7E**). This result suggests that WormClust can be used to ‘deorphan’ unannotated metabolic genes. As another example, we found that *acdh-11* is highly coexpressed with mitochondrial fatty acid degradation and isoleucine degradation genes (**Fig 7F**). Therefore, we propose that *acdh-11* can now be added to the iCEL model as another OR gene to reactions in these pathways. We envision future systematic studies of orphan metabolic genes and TFs to increase the annotation of metabolic genes and elucidate the transcriptional mechanisms that regulate their expression.

## Discussion

In this study, we performed a systems-level analysis of mRNA level variation as a proxy for the transcriptional regulation in *C. elegans*. Even with our conservative approach, we found that most metabolic genes are under transcriptional control. By a combination of supervised and unsupervised methods, we found that genes in many metabolic pathways are coexpressed and identified gene clusters that represent parts of pathways, or branches of pathways with strong coexpression.

What is the purpose of transcriptional activation or repression of metabolic genes? We propose that the transcriptional regulation of metabolic genes can serve different purposes. First, there is extensive transcriptional regulation of metabolic genes in different tissues. This can be viewed as the setup of metabolic network functionality depending on a tissue’s needs. Indeed, tissues have different needs. For instance, the *C. elegans* intestine serves as the entry point of bacterial nutrients and requires the expression of enzymes that aide digestion and metabolite transport to other tissues. Similarly, the animal’s muscle needs to produce energy to support movement and is therefore highly catabolic. Second, metabolic genes are transcriptionally regulated during development. This likely reflects the need for different aspects of metabolism as tissues differentiate and grow and as different metabolic functions are necessary. We refer to the expression of different metabolic genes and pathways in different tissues and different developmental stages as ‘metabolic network wiring’. A third function of metabolic gene activation or repression is under different conditions that dictate the need for different metabolic functions. This can be for the breakdown of different nutrients, e.g., carbohydrates versus fats, versus protein, or to rewire metabolic pathways when others are perturbed. An example of this is the propionate shunt, which is transcriptionally activated when flux through the preferred, vitamin B12-dependent pathway is perturbed (Bulcha et al., 2019; Watson et al., 2016). We refer to the transcriptional rerouting of metabolism as ‘metabolic rewiring’. Finally, metabolic genes can be transcriptionally activated when flux through the pathway in which they function is hampered. An example of this is the Met/SAM cycle in *C. elegans*, which is transcriptionally activated when flux through the cycle is low, for instance under low vitamin B12 dietary conditions (Giese et al., 2020). For genes encoding enzymes that function in multiple reactions and metabolic pathways (e.g., *alh-8*), we can learn with which pathways they are more strongly coexpressed. Our study provides a facile portal to investigate the tissues, developmental stages, or conditions under which particular metabolic genes and pathways are highly coexpressed, which will help to formulate hypotheses for detailed follow-up studies.

Genes that are coexpressed often function together and are frequently coexpressed with their transcriptional regulators (Eisen et al., 1998; Hughes et al., 2000; Kim et al., 2001; Segal et al., 2003; Stuart et al., 2003). We used this principle to develop WormClust with which any *C. elegans* gene, including TFs, can be used to search for metabolic pathways with which it is coexpressed. However, it is important to note that the most critical regulators, those that respond to the initial information, are often not coexpressed with their target genes. Indeed, *nhr-10*, which is essential for activation of the propionate shunt in response to high levels of propionate, does not change much in expression under relevant conditions (Bulcha et al., 2019). To identify such ‘first responders’, it will be useful to employ promoter-reporter strains with large scale RNAi screens (Bhattacharya et al., 2022; MacNeil et al., 2015). Further, at least half of all *C. elegans* metabolic genes are not yet associated with reactions or pathways (Yilmaz et al., 2020). We propose that such genes may be ‘deorphaned’ in the future using WormClust. Longer term, we envision that association of other types of regulators, such as RNA binding proteins and microRNAs, with metabolic pathways can be used to gain broader insights into the functional connections among different biological processes.

Finally, the approaches used here should be broadly applicable to any organism, including humans, for which large gene expression profile compendia and high-quality metabolic network models are available. By applying these approaches, deeper insights into the transcriptional control of metabolism will be obtained, as well as insights into the conditions under which metabolic genes and pathways are activated or repressed.

## Materials AND Methods

### Preprocessing of Genes

The master list of *C. elegans* genes considered for analysis were downloaded from WormBase public ftp site (release WS282) (Harris et al., 2020). The genes were filtered out to obtain only live and protein-coding genes, which amounted to 19,985 genes in total.

### Development Dataset

Post-embryonic expression profiles were based on published RNA-seq data (Kim et al., 2013). Briefly, the authors measured the transcriptome of wild-type (N2) animals from hatching to 48h post-hatching every two hours. This dataset includes 21,714 protein-coding genes including 2,405 metabolic genes. Genes were classified into 12 clusters based on their expression profiles (Kim et al., 2013). We refer to the cluster showing relatively invariant expression as the “flat cluster” and the genes within this cluster as “flat genes”. Selecting only live protein-coding genes (WS282) resulted in a total of 18,113 genes, including 2,397 metabolic genes. Of these, 4,689 are flat genes, including 995 metabolic genes. We generated a histogram of average gene expression across development using the logarithm of reads per kilobase per transcript (RPKM) values at base 2. This resulted in a bimodal expression distribution that was fitted by two superposed Gaussian curves, representing a high expression subpopulation and a low expression subpopulation (LES). Genes that showed expression values less than the mean plus the standard deviation of LES at all the time points were filtered out to avoid false fluctuations in gene expression. After this step, a total of 14,561 genes were left, including 2,184 metabolic genes. The number of flat genes was reduced to 4,646, including 986 metabolic genes.

### Tissue Dataset

Tissue-level expression profiles were based on a single-cell RNA-seq dataset of animals at the second larval stage (L2) (Cao et al., 2017). This dataset provides gene expression as transcripts per million (TPM) for 20,271 protein-coding genes including 2,506 metabolic genes across seven major tissues: body wall muscle, glia, gonad, hypodermis, intestine, neurons, and pharynx. Selecting only live genes (WS282) reduced the number of genes to 19,675 including 2,491 metabolic genes. The dataset was previously processed to label gene expression in every tissue according to the level of expression into four categories: high, moderate, low, and rare (Yilmaz et al., 2020). Genes that showed rare or low expression in all seven tissues were filtered out in this study, resulting in 13,305 genes including 2,143 metabolic genes.

### Gene Expression Compendium

A compendium of gene expression datasets was generated using a combination of public datasets. First, 374 microarray, RNA-Seq and tiling array datasets related to *C. elegans* were downloaded from WormBase (Harris et al., 2020). Then, only those datasets that consisted of at least ten conditions were selected, resulting in 169 datasets. These datasets were individually examined for batch effects, since many were obtained from multiple microarray experiments where total RNA was not normalized. Initially, histograms of expression values were analyzed, and sixteen microarray datasets that displayed abnormal distributions where the correlation distributions were skewed towards +1, hence suggesting that the data may consist of samples that are highly distinctive from each other or are from separate experiments altogether. Such datasets were selected for further examination (**Fig EV3A**). Twelve of these datasets were found to be composed of two subsets of data, where all genes in one subset were up- or down-regulated with respect to the other except for a few. The samples forming each subset were independent of the other and seemed to have different amounts of total RNA or a similar batch effect. Therefore, these datasets were divided into two separate datasets to correct for batch effects. The remaining four datasets were removed since the source of abnormalities in their distributions of expression was not clear. This processing resulted in a total of 177 datasets. For each dataset, the expression of every gene was normalized by converting the expression values to z-scores based on expression across all conditions using the Normalizer function of Sleipnir library (Huttenhower et al., 2008). Once all the datasets were z-normalized, they were combined to form a compendium with 4,796 conditions (sum of multiple conditions within 177 datasets) using the Combiner function of the Sleipnir library (Huttenhower et al., 2008), which took a union set of all genes across the different datasets and converted missing values to NaN (not a number) for subsequent processes.

### Calculation of Variation Score In the Development Dataset

We define Variation Score (VS) as a measure of the deviation of a gene’s expression profile from a flat reference over time in the development dataset (Kim et al., 2013). Prior to any analysis, expression values of every gene were normalized by total expression in all time points using equation (1),

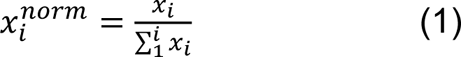

where x_i_ indicates the expression value of gene x at time i. To define a reference profile of invariant expression, a line was constructed in time by joining the mean normalized expression value of the flat cluster at every time point. An envelope around this line was then defined by adding and subtracting the standard deviation of each point. A deviation from this envelope, referred as Variation Score (VS), was then computed by taking the average distance between an individual gene profile and the reference profile according to Eq.2,

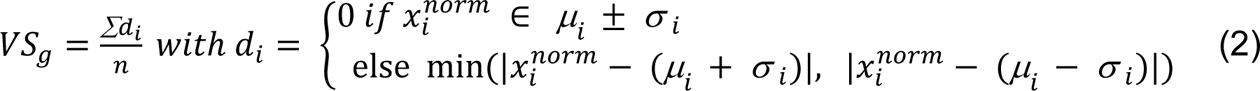

where n is the number of observations for the gene g, d_i_ is the distance at time i between the normalized level of expression of the gene *s_i_^snorm^* and the closest border of the reference flat profile, and μ_i_ and σ_i_ are mean and standard deviations of normalized expression values of flat genes at time point *i* respectively. A graphical example of this calculation is provided in **Fig EV1A**. With this definition, a VS = 0 means that the profile of a given gene stays within the envelope of the flat cluster and is therefore perfectly flat, or invariant.

### Calculation of Coefficient of Variation

Coefficient of Variation (CV) is a statistical measure that is used to calculate the dispersion of data. For every gene, CV was calculated by dividing standard deviation of expression across different samples (σ) (*e.g.*, different tissues in case of tissue dataset) to the mean of expression across samples (μ). CV was empirically thresholded using the CV of known propionate shunt genes to keep the approach conservative.

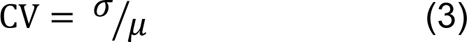

### Calculation of Coexpression of Gene Pairs

The correlation in expression of metabolic gene pairs during development, across tissues, and across the compendium of gene expression studies were calculated based on Pearson Correlation Coefficient (PCC) using the Distancer function of Sleipnir library (Huttenhower et al., 2008). These correlations defined pairwise coexpression. Differences in the distribution of coexpression values between random, AND genes, OR genes, other paralogs, all pathway genes and PW genes were evaluated using Mann-Whitney U test (Fay and Proschan, 2010).

### Custom Pathway Enrichment Analysis Pipeline

Pathway-to-gene annotations from level 4 of WormPaths (Walker et al., 2021) were used as input gene sets. Each metabolic pathway (or category such as an enzyme complex; hereafter referred to as pathway for simplicity) consists of two or more annotated metabolic genes. The coexpression of genes in each metabolic pathway with all other genes in the metabolic network was extracted. Subsequently, the mean of the correlations of pathway genes with all other metabolic genes (excluding the self-correlations) was calculated. The mean values were used to define a ranked list of metabolic genes for every metabolic pathway. GSEA was then performed on the pre-ranked list of each pathway using the PreRank module (Subramanian et al., 2005). Enrichment score (ES) is the degree to which the genes in a gene set are overrepresented at the top or bottom of the entire ranked list of genes. Since the genes that are functionally related are mostly positively correlated, we only consider the genes at the top of the list, hence the ones positively contributing to ES. Leading edge subset enlists the gene hits before the peak while calculating ES, therefore consisting of genes that contribute the most to the enrichment score. NES was derived for each pathway by normalizing the ES values to mean ES for all permutations of the gene sets. This accounts for differences in gene-set size. FDR is the estimated probability that a gene set with a given NES represents a false positive finding. The significance cutoff for the GSEA was set at an FDR value of ≤ or equal to 0.05. If a pathway was found to be significantly self-enriched, it was categorized as coexpressed.

### Unsupervised Approach That Combines Coflux and Coexpression

The first part of the approach involves using flux balance analysis (FBA) to simulate reaction rates (fluxes) in the metabolic network and then using a flux dependency metric, referred to as coflux, to measure pairwise associations of genes in the *C. elegans* metabolic network model (Yilmaz et al., 2020; Yilmaz and Walhout, 2016). We examined all reaction pairs to see if constraining the flux of one of the reactions to zero reduces the flux of the other (see below for the algorithm). The coflux value is zero for independent reactions and one for reactions that are fully coupled. Reactions that are connected by a junction to another reaction are usually partially dependent. After generating coflux values for each pair of reactions, we converted the reaction matrix to a pairwise gene coflux matrix using GPR associations. A high coflux value for a gene pair indicates that the genes encode enzymes acting in the same metabolic process. For the second part of the approach, a coexpression matrix was derived from the *C. elegans* gene expression compendium described above. All negative correlations were converted to zero to be consistent with the coflux matrix. The coflux and coexpression matrices were multiplied to obtain a product matrix. Since both coexpression and coflux values are between 0 and 1, a product takes a high value only if both coflux and coexpression values are high. Hierarchical clustering was then performed on the product matrix using dynamic cut tree algorithm using cutreehybrid package (Langfelder et al., 2008).

### Coflux Algorithm

The coflux value for each gene pair was calculated using FBA with iCEL1314 (Yilmaz et al., 2020). First, the standard bacterial diet was amended with a minimum set of nutrients (*i.e.*, by allowing uptake through exchange reactions in the model as indicated in **Table S8**) that warranted non-zero flux in all reactions of the model. Then the following steps were taken to calculate coflux values:

- For every irreversible reaction *i*,

- Calculate v_max,*i*_, the maximum flux that can be achieved with the intact network.
- For every reaction *j*, calculate v_max,*ij*_, which is the maximum flux observed in reaction *i* when reaction *j* is constrained to a flux of zero. If *i* is included in a predefined set of 15 redundant reaction pairs (*i.e.*, reactions with similar reactants and products except for differences such as the use of NADP instead of NAD as electron carrier, **Table S8**), the flux of the corresponding reaction in the pair was also constrained to zero.
- For every reaction *j*, calculate the coflux with *i (c_ij_)* using Eqn 4.

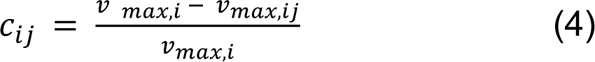
- For every reversible reaction i,

- For every reaction *j,* repeat the above steps to calculate the coflux with i in forward direction (*c_ij, forward_*).
- Calculate v_min,*i,*_ the minimum (*i.e*., the most negative, as negative flux indicates flux in reverse direction) flux that can be achieved with the intact network.
- For every reaction *j*, calculate v_min,*ij*_, which is the minimum flux observed in reaction *i* when reaction *j* is constrained to a flux of zero. Once again, the reaction redundant with reaction *i* is also constrained to zero flux, if applicable.
- For every reaction *j*, calculate the coflux with the reverse direction of *i* (*c_ij,reverse_*) using Eqn 5.

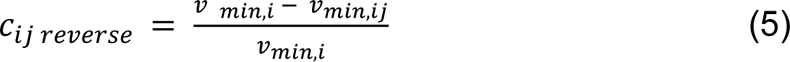
- For every reaction j, calculate final coflux with i as the maximum of *c_ij, forward_* and *c_ij, reverse_* (Eqn. 6).

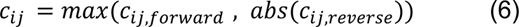
- Since *c_ij_* and *c_ji_* are not necessarily equal, calculate final coflux value for every reaction pair using Eqn. 7.

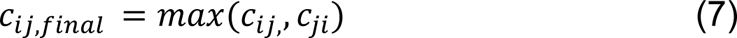
- Convert the reaction coflux matrix to a gene coflux matrix based on gene-reaction associations. If a gene pair is associated through multiple reaction pairs (*i.e.*, when at least one of the genes is associated with multiple reactions), take the maximum of coflux values between reactions to calculate gene coflux.

### Hierarchical Clustering

Hierarchical clustering was performed using average method of linkage on the dissimilarity matrix generated by 1 minus the product matrix value (coflux*coexpression). Dynamic cut tree algorithm from cutreehybrid package was used to cut the dendrogram generated by this clustering with stringent parameters deepSplit=2 and minClusterSize=3 and relatively relaxed parameters deepSplit=3 and minClusterSize=6 (Langfelder et al., 2008). The stringent setting was thresholded based on the occurrence of propionate shunt genes together in one cluster while keeping the size of the smallest cluster to be at least 3. The relaxed setting was chosen to capture larger clusters, such as the Met/SAM cycle genes in a single cluster.

### Quantifying Cluster Quality Through Silhouette Score

To assess the quality of clustering, we first calculated silhouette score of each metabolic gene based on its placement in each cluster and then calculated MSS for every cluster. Silhouette score determines the quality of clustering by measuring the cohesiveness of genes within the same cluster and separateness from the genes in the neighboring clusters (Rousseeuw, 1987). It was calculated using scikit-learn package (Pedregosa, 2011).

### Finding Activation and Repression Conditions of Metabolic Clusters

To find activation and repression conditions of each cluster, we first calculated the mean coexpression of all gene pairs in that cluster in each dataset separately. To rank datasets that showed highest coexpression uniquely for each cluster, we normalized these mean coexpression values of all clusters using z-scoring across each dataset. We then identified the 30 best datasets that potentially represents activation/repression conditions of each cluster by the z-score values of mean coexpression. After this, we manually inspected each dataset in the order of decreasing mean coexpression, its associated published paper, and the heatmap to understand activation/repression conditions.

### Gene-Centric Coexpression with Metabolic (Sub)-Pathways by WormClust

We developed a custom computational pipeline that identifies coexpression of *C. elegans* genes with metabolic genes used in this study. The pathway-gene sets were generated using WormPaths as a GMT (Gene Matrix Transposed) file, a tab-delimited file of gene sets (Walker et al., 2021). The ranked coexpression list of metabolic genes was extracted for each queried gene, from the global coexpression matrix generated using compendium of 177 datasets. The ranked list of each queried gene was used to run Gene Set Enrichment Analysis (GSEA) on the custom metabolic pathway-gene sets.

### Data and Code Availability

Gene clusters from unsupervised approach and pathway enrichment of all protein coding genes outside the iCEL model, including but not limited to TFs, orphan metabolic genes, and transporters are available using WormClust on the WormFlux website (wormflux.umassmed.edu). Other data can be found in supplementary tables. We also created a Github repository (https://github.com/WalhoutLab/WormClust) for this project, which includes scripts that generated results presented here.

## Acknowledgements

We thank members of the Walhout lab and Caryn Navarro, for discussion and critical reading of the manuscript. This work was supported by National Institute of Health (NIH) grant R35GM122502 to A.J.M.W and R01HG005084 to C.L.M.

## Author Contributions

Conceptualization, S.N. and A.J.M.W; Methodology, S.N, M.A.J. and L.S.Y; Software, S.N. and L.S.Y.; Formal analysis, S.N.; Investigation, S.N.; Writing-Original draft, S.N. and A.J.M.W.; Writing-Review & Editing, S.N, W.W., M.A.J., C.L.M., L.S.Y., and A.J.M.W.; Visualization, S.N.; Supervision, C.L.M., L.S.Y., and A.J.M.W.; Funding acquisition, A.J.M.W.

## Expanded View Figure Legends

**Figure EV1.**
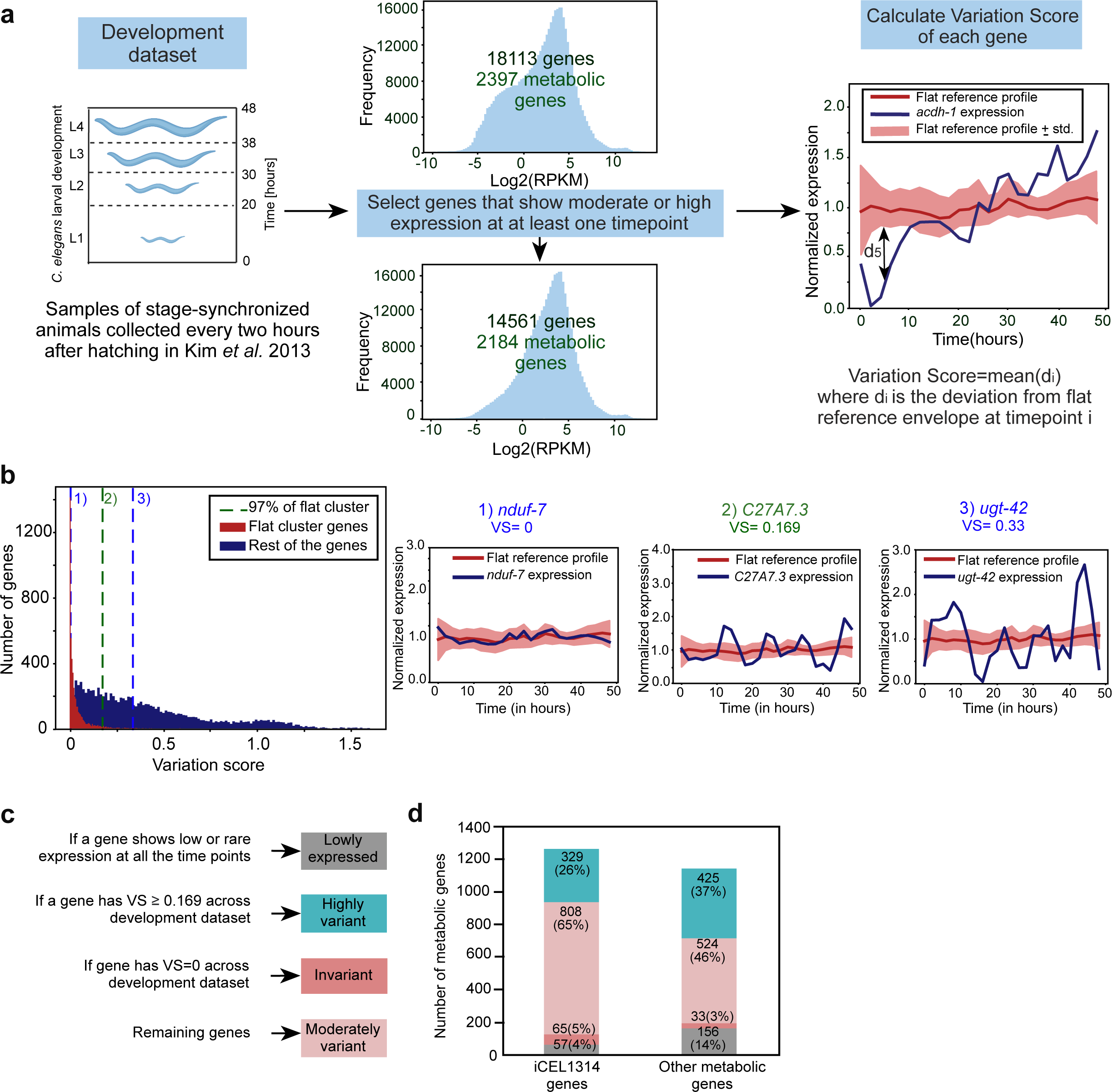
Transcriptional Regulation of Metabolism during Development. (A) Computational pipeline to identify highly variant metabolic genes during development. Genes that showed either moderate or high expression in at least one time point were selected, reducing the number of genes from 18,113 (2,397 metabolic) to 14,561 (2,184 metabolic). For each gene, VS was calculated using the deviation from a flat reference profile at each time point (see **Methods**). The red line indicates the mean value and light red shaded area the standard deviation of the flat reference profile. The profile of *acdh-1* is plotted in blue as an example of a developmentally regulated gene. d_5_ denotes the deviation of *acdh-1* expression from the flat reference profile at the 5^th^ data point. (B) Distribution of VS of genes belonging to the flat cluster versus those belonging to other clusters. The vertical lines at 1 and 3 represent the iCEL1314 genes *nduf-7* and *ugt-42,* that have the lowest and highest VS of the flat set, respectively. The vertical line at 2 indicates the gene *C27A7.3* with selected threshold of VS at the 97% quantile of the flat cluster (VS = 0.169). (C) Diagram showing criterion of categorizing metabolic and non-metabolic genes into four categories across development: lowly expressed, invariant, moderately variant and highly variant. (D) Bar graph shows the distinction of low expressed, invariant, moderately variant and highly variant genes during development in iCEL1314 and other metabolic genes. Color legend as indicated in (C).

**Figure EV2.**
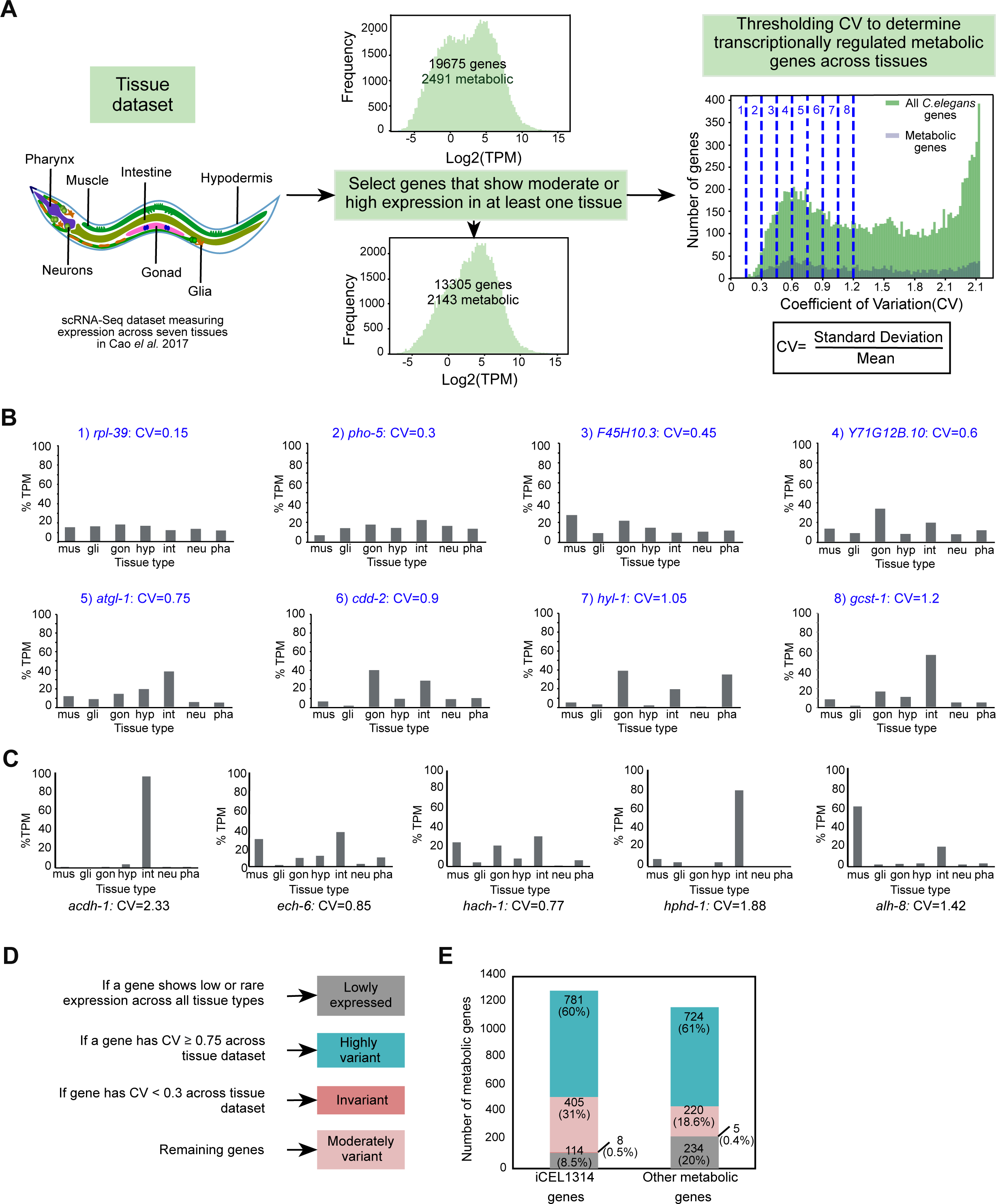
Transcriptional Regulation of Metabolic Genes across Different Tissues. (A) Computational pipeline to determine highly variant metabolic genes across tissues. Genes that show either moderate or high expression in at least one tissue are selected for analysis reducing the number of *C. elegans* genes from 19,675 (2,491 metabolic) to 13,305 (2,143 metabolic). The coefficient of variation (CV) of each gene was calculated by dividing the standard deviation of expression across tissues by the mean expression. Different thresholds of CV were titrated to select a stringent CV to categorize genes as variant and therefore potentially transcriptionally regulated. Examples for each threshold are provided in (B). (B) Bar graphs showing expression profiles of example genes across tissues with CV 0.15, 0.3, 0.45, 0.6, 0.75, 0.9, 1.05 and 1.2. Numbering of examples is according to the corresponding threshold lines in (A). (C) Tissue expression and CV values of propionate shunt genes. (D) Diagram showing criterion of categorizing metabolic and non-metabolic genes into four categories across tissues: lowly expressed, invariant, moderately variant and highly variant. (E) Bar graph shows the distinction of low expressed, invariant, moderately variant and highly variant genes across tissues in iCEL1314 and other metabolic genes. Color legend as indicated in (D).

**Figure EV3.**
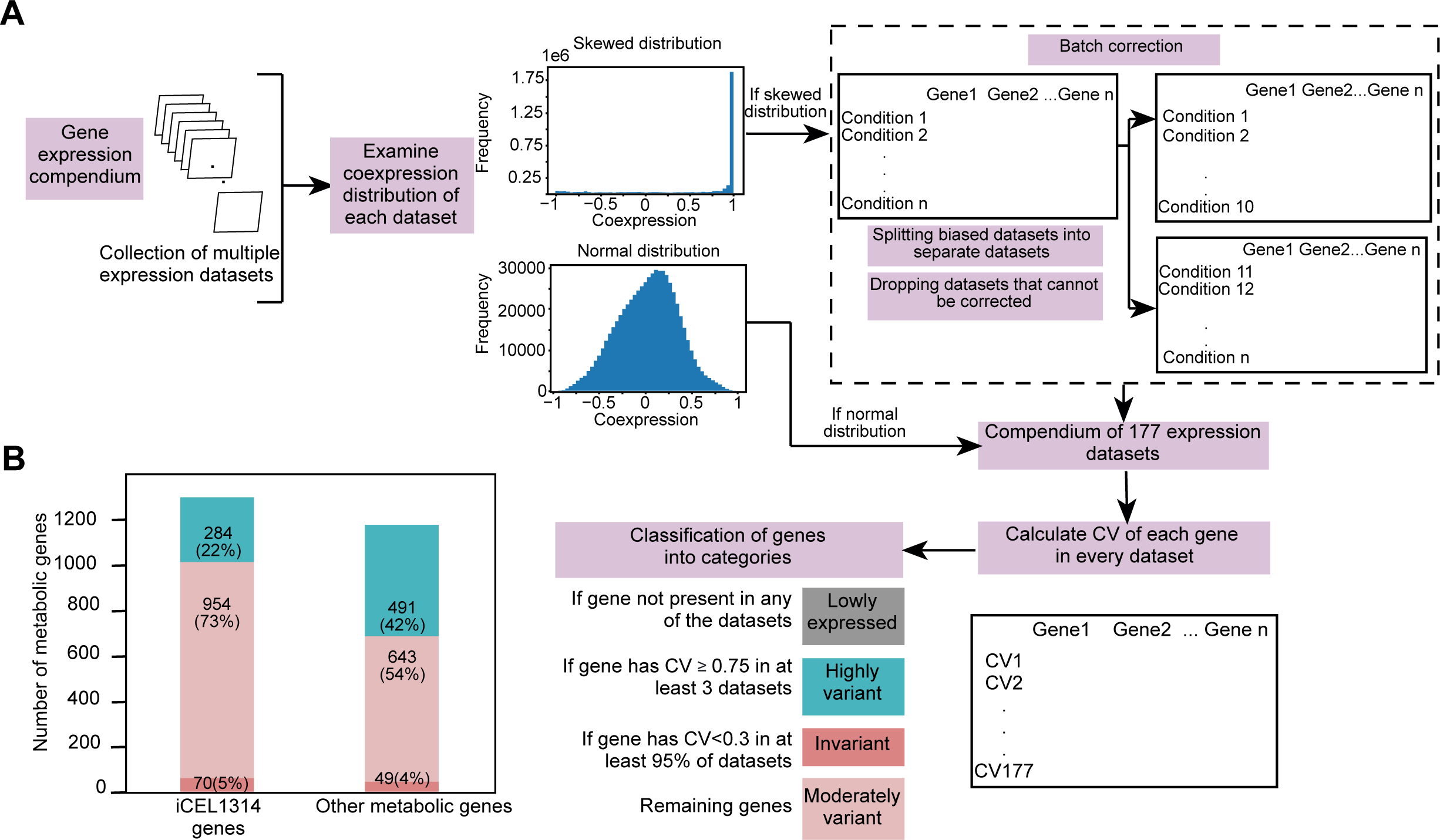
Transcriptional Regulation of Metabolic Genes across a Compendium. (A) Computational pipeline to determine transcriptionally regulated metabolic genes across the gene expression compendium. Datasets were evaluated for batch effects using correlation distribution. Some skewed datasets were corrected for batch-effect by splitting into separate datasets, while some were removed if the source of skewness was not clear. This resulted in 177 datasets for analyses. CV of each gene was calculated. The criterion of classifying genes is based on the number of datasets with high CV and fraction of datasets with low CV. (B) Bar graph showing low expressed, invariant, moderately variant and highly variant genes in the compendium in iCEL1314 and other metabolic genes. Color legend as indicated in (A).

**Figure EV4.**
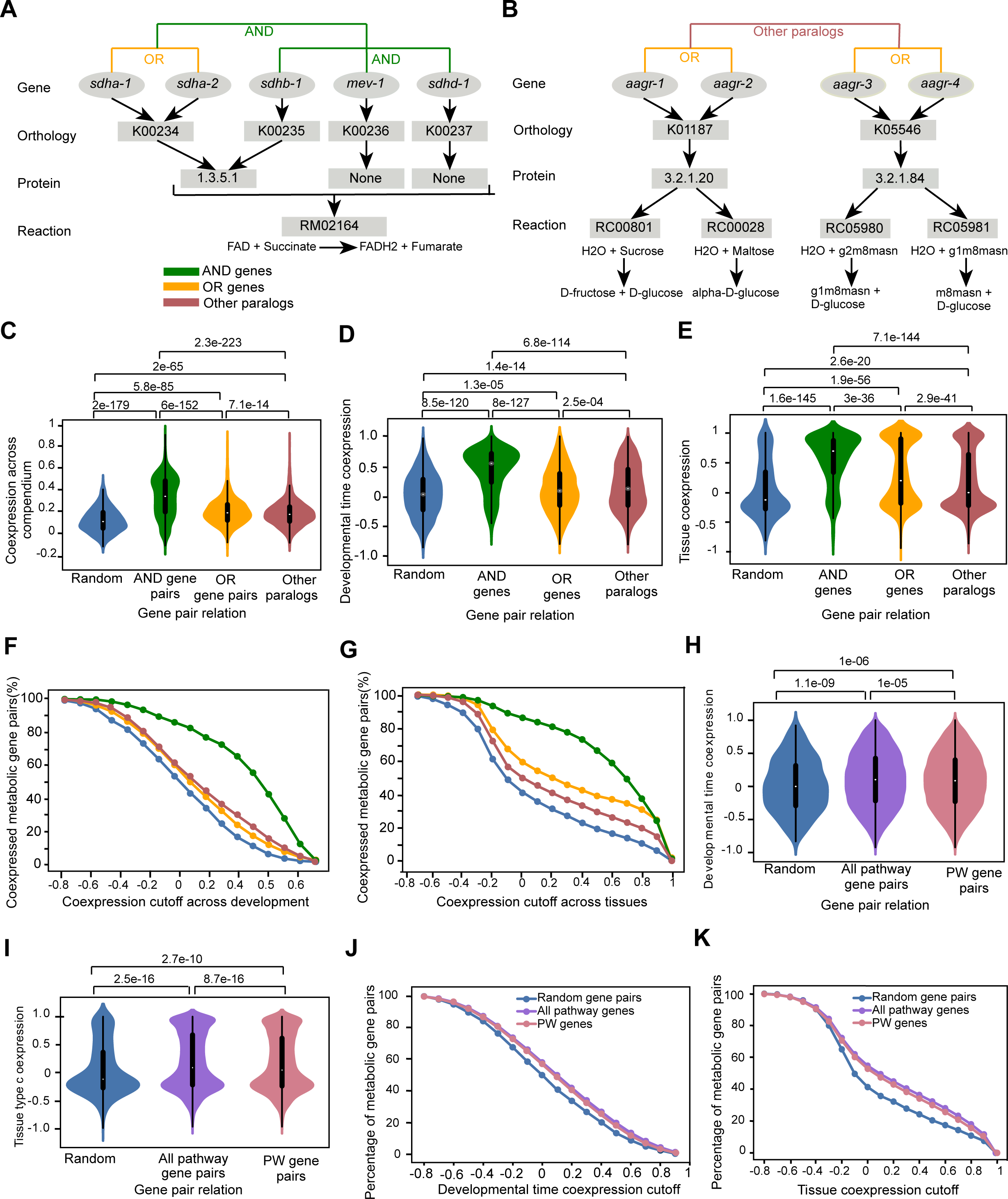
Reaction-Level Analysis of Metabolic Pathways. (A) The conversion of succinate to fumarate, which is part of complex II of the ETC and of the TCA cycle, is carried out by succinate dehydrogenase. Diagram showing that succinate dehydrogenase is composed of the OR genes *sdha-1* and *sdha-2* that each function together with the rest of the genes as AND genes. The GPR of this reaction is noted as [(*sdha-1* | *sdha-2*) & *sdhb-1*] & *mev-1* & *sdhd-1*]. (B) Example of a gene family (*aagr*) where members occur as paralogous OR gene pairs in separate reactions. Pairs of paralogs associated with different reactions are called other paralogs. (C-E) Violin plot comparing coexpression for different populations of gene pairs including random, AND, OR and other paralogs gene pairs in compendium of expression datasets (C), across development (D), and across tissues (E). (F-G) Percentages of AND, OR, other paralogs and random metabolic gene pairs categorized as coexpressed using different coexpression values as cutoffs across developmental time (F) and tissues (G). (H-I) Violin plots showing coexpression of different populations of gene pairs across different developmental stages (H) and tissues (I). (J-K) Percentage of different types of gene pairs categorized as coexpressed using different coexpression values as cutoffs across developmental time (J) and tissues (K).

**Figure EV5.**
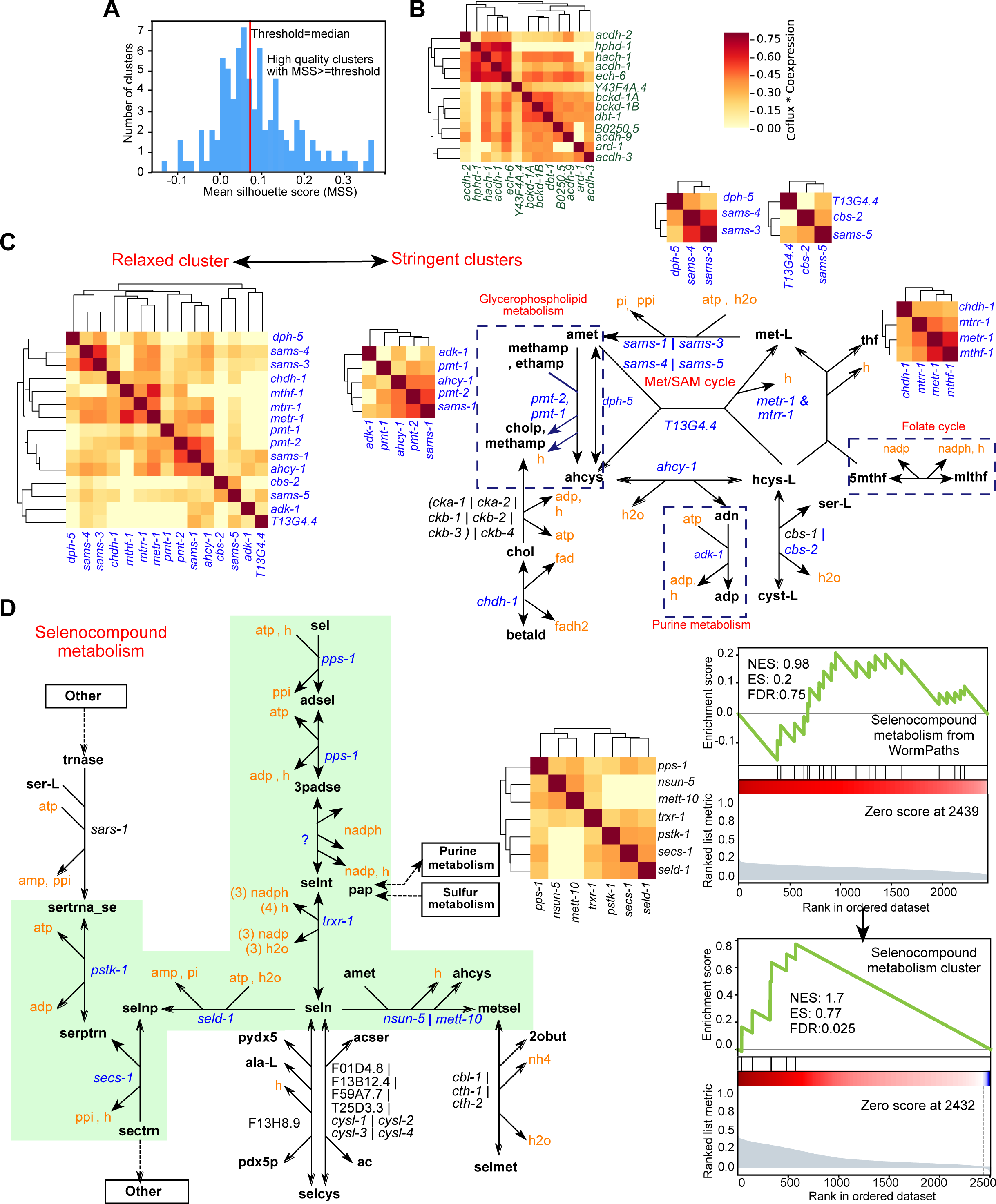
Unsupervised Clustering of Product Matrix using Relaxed Parameters. (A) Plot showing the distribution of the MSS of all clusters obtained using relatively relaxed parameters (deepSplit=3, minClusterSize=6). The threshold to define high-quality clusters and examine potential coexpressed metabolic units is chosen as the median of MSS. (B) Distinct cluster of known coregulated metabolic pathway propionate shunt genes together with isoleucine and valine degradation genes obtained using relaxed clustering parameters (deepSplit=3, minClusterSize=6) with the dynamic cut tree algorithm. (C) Clusters of Met/SAM cycle genes and genes of related pathways obtained using relaxed (deepSplit=3, minClusterSize=6) and stringent (deepSplit=2, minClusterSize=3) parameters. Stringent clusters are shown near the drawn pathways of clustered genes. (D) Pathway boundary (green shade) within selenocompound metabolism defined by genes in a high-quality cluster obtained by relaxed parameters and shown by the clustered heatmap (left). Mountain plots showing comparison of self-enrichment analysis statistics (NES, ES and FDR) of selenocompound metabolism from WormPaths with that of selenocompound cluster obtained by the unsupervised approach (right).

